# The embryo-derived protein PDI is highly conserved among placental mammals and alters the function of the endometrium in species with different implantation strategies

**DOI:** 10.1101/2024.05.02.592140

**Authors:** Haidee Tinning, Alysha Taylor, Dapeng Wang, Anna Pullinger, Georgios Oikonomou, Miguel A. Velazquez, Paul Thompson, Achim Treumann, Peter T. Ruane, Mary J O’Connell, Niamh Forde

## Abstract

Pregnancy establishment in mammals requires a complex sequence of events, including bi-lateral embryo-maternal communication, leading up to implantation. This is the time when most pregnancy loss occurs in mammals (including humans and food production species) and dysregulation in embryo-maternal communication contributes to pregnancy loss. Embryo-derived factors modify the function of the endometrium for pregnancy success. We hypothesise that these previously unexplored conceptus-derived proteins may be involved in altering the function of the endometrium to facilitate early pregnancy events in mammals with different early pregnancy phenotypes. Here, we show that protein disulphide-isomerase (PDI) is a highly conserved protein among mammals, and provide evidence for a species-specific roles for PDI in endometrial function in mammals with different implantation strategies. We show how PDI alters the endometrial transcriptome in human and bovine *in vitro* in a species-specific manner, and using a microfluidic approach we demonstrate that it alters the secretome capability of the endometrium. We also provide evidence from *in vitro* assays using human-derived cells that *MNS1,* a transcript commonly downregulated in response to PDI in human and bovine endometrial epithelial cells, may be involved in the attachment (but not invasion) phase of implantation. We propose that the trophoblast-derived protein PDI, is involved in supporting the modulation of the uterine luminal fluid secreted by the endometrium to support conceptus nourishment, and also in the process of embryo attachment to the uterine lumen for pregnancy success in mammals.

**SIGNIFICANCE STATEMENT:** We provide evidence that a highly conserved protein (PDI) alters the endometrial transcriptome in a species- and cell-specific manner. Exposure of endometrial epithelia to PDI altered genes belonging to immune modulatory, pro-inflammatory, and adhesion-pathways. One transcript, MNS1, was commonly downregulated in endometrial epithelia from species with superficial (bovine) and invasive (human) implantation morphologies. Knockdown of MNS1 expression in humans epithelia altered the ability of human trophoblast BeWo spheroids to attach suggesting a mechanism by which PDI affects implantation in human and bovine. In addition, using a microfluidics approach we have shown that PDI alters the secretome in a species-specific manner demonstrating PDI alters a key function of the endometrium in mammals.

## INTRODUCTION

Placental mammals display a diversity in the timing, morphology, and molecular cues required for successful pregnancy (1). Despite this diversity, the majority of pregnancy loss occurs in the pre- and peri-implantation stages of pregnancy in most mammalian species studied (2). While some of these losses are attributable to defects in gametes and embryo development, transfer of competent embryos does not always result in pregnancy success. Dysregulation in signalling or communication between developing embryo and endometrium contributes to pregnancy loss.

The endometrium is a specialised heterogeneous tissue which lines the uterine cavity. Its main functions are to support growth and development of the embryo, respond to the pregnancy recognition signals from the embryo/conceptus, and facilitate receptivity to implantation. These functions are mediated by the different heterogeneous endometrial cells including fibroblast-like stromal cells, immune cells, a microvasculature system, and specialised epithelial cells. The endometrium also contains secretory glands which, along with the luminal epithelium, secrete the uterine luminal fluid (ULF). Secretion of the ULF is necessary to support growth and development of the embryo prior to the formation of the placenta (3). The endometrium responds spatiotemporally to steroid hormones in the maternal circulation, including progesterone, concentrations of which rise and fall in a cyclic manner during the estrus or menstrual cycle. Progesterone-induced changes in the endometrium are essential for processes such as establishing receptivity to implantation. The ULF also changes in composition in response to progesterone (4, 5), and in response to signals from the embryo/conceptus (6).

Previous studies have determined that the transcriptomic response of the endometrium to the bovine conceptus during the pregnancy recognition period is greater than that of pregnancy recognition signal alone (7). To determine what other proteins may be involved in altering the endometrial transcriptome/ endometrial function, Day 16 conceptus conditioned medium was subjected to quantitative proteomic analysis. Amongst the most abundant proteins was Protein disulphide-isomerase (PDI) (also known as protein disulphide-isomerase precursor or prolyl 4- hydroxylase β-subunit [P4HB]) (8). PDI is a 508 amino acid polypeptide belonging to the thioredoxin family of proteins and has multiple cellular functions, broadly involved in 1) redox regulation, 2) disulphide isomerase activity, 3) chaperone protein activity, and 4) redox-dependent chaperoning (9). It is a highly abundant cellular protein, and has been found to be upregulated in different cancers (10) where is it often associated with poor outcomes (11), progression and invasion (12). PDI has also been linked with protein-misfolding associated neurodegenerative disorders, some cardiovascular disorders, and pathogen cellular entry in some infectious diseases such as HIV (13).

With several known functions, and the capacity to act as a sub-unit in multiple known complexes, PDI is involved in many cellular processes. In the endoplasmic reticulum, PDI is involved in the maintenance (forming and breaking) of disulphide bonds between cysteine residues during protein folding (14). PDI can also stabilise and destabilise disulphide bonds in MHC class I molecules and is therefore involved in the ability of a cell to present antigens to the immune system (15). PDI can also act as a chaperone protein for misfolded proteins in the endoplasmic reticulum (16). PDI can also have other roles when it associates with other proteins as a complex, e.g., PDI is the β-subunit of microsomal triacylglycerol transfer protein (MTP) which is involved in incorporating triacyclglycerols into lipoproteins. PDI can also act as the 2 β-subunits of prolyl 4-hydroxylase (P4H), which catalyses procollagen pro-α-chain proxlyl hydroxylation in the endoplasmic reticulum (17).

More recently, it was proposed that PDI is also involved in endometrial receptivity to implantation in humans (18). Fernando *et al* (2021) found that oestrogen and progesterone altered the expression of certain PDI isoforms in different human endometrial epithelial cell lines (18). Inhibition of PDI activity increased the attachment of Jeg-3 spheroids to AN3CA endometrial epithelial cells. Oestrogen treatment has been shown to upregulate PDI expression in the endometrium of mice (19), and PDI expression is higher in the mid-secretory phase compared to the early-secretory phase endometrium of humans with unexplained infertility (20). We have also shown that PDI alters expression of a core set of microRNAs in endometrial epithelia demonstrating its potential as a regulator of endometrial function (21). PDI is a highly conserved conceptus-derived protein produced at the time of pregnancy recognition in cattle (18). We sought to determine if the role of PDI in implantation, including its effect on maternal endometrial tissues, is conserved among mammals with differing early pregnancy processes.

## MATERIALS AND METHODS

Unless otherwise stated all materials were sourced from Sigma-Aldrich.

### Conservation analysis of PDI

Analysis of species conservation was carried out as previously described (22). Briefly, orthologs of PDI across seventeen species from a range of placental mammals, non-placental mammals, and non-mammal outgroups were identified and extracted from Ensembl (23). Multiple sequence alignment was performed using MAFFTv7.3 (24) under default settings, and using SeaView (25) for visualisation. The percentage identity matrix was generated in Clustal Omega (26).

### Recombinant PDI production

Recombinant bovine PDI (rbPDI) was produced at the Newcastle University Protein and Proteome Analysis centre using an *E. coli* pET3A expression vector system and subsequent purification steps, as described in (22). Recombinant ovine IFNT (roIFNT) was kindly gifted by Professor Fuller Bazer for this study, produced in *P.pistoris* to >80% purity as described by Van Heeke *et al* (27). All recombinant proteins were eluted/purified into PBS and stored at -80°C prior to use.

### Primary bovine endometrial cell culture

Primary bovine endometrial epithelial cells (bEECs) and primary bovine endometrial stromal cells (bESCs) were isolated by enzymatic digestion from late-luteal phase uterine tracts (n=3), as described previously (22). bEECs and bESCs were maintained in complete bovine medium (RPMI 1640 [Gibco, Massachusetts, US], 10% charcoal-stripped FBS [PAA Cell Culture Company, UK], 1% antibiotic antimycotic solution [ABAM, Sigma-Aldrich, Missouri, USA]). bESCs at a concentration of 150,000 cells per well (n=5) and bEECs at a concentration of 300,000 cells per well (n=5) were plated into separate 6-well plates in 2 mL complete bovine medium. All cultures were maintained at 37°C/5% CO_2_ in a humidified incubator. At 70% confluence bEECs and bESCs were treated with one of the following for 24 hr: (1) Media control, (2) Vehicle control (PBS only), (3) 10 ng/ml rbPDI, (4) 100 ng/ml rbPDI, (5) 1000 ng/ml rbPDI or, (6) 1000 ng/ml rbPDI combined with 100 ng/ml roIFNT.

### Human endometrial cell culture

Ishikawa (ECACC 99040201, passage 10) immortalised human endometrial epithelial cells (hEECs) were plated in biological triplicate at 300,000 cells per well in a 6 well plate, in 2 mL complete human medium (DMEM/F12 [Gibco, Massachusetts, US], 10% FBS [PAA Cell Culture Company, UK], 1% GSP [Gibco, Massachusetts, US]). All cultures were maintained at 37 °C/5% CO_2_ in a humidified incubator. At 70% confluence hEECs were treated with one of the following for 24 hours: (1) Media control, (2) Vehicle control (PBS only), (3) 10 ng/ml rbPDI, (4) 100 ng/ml rbPDI, or (5) 1000 ng/ml rbPDI.

### RNA extraction, cDNA conversion, and qRT-PCR

Following treatment cells were washed with PBS, lysed in 400µL *mir*Vana lysis solution from *mir*Vana RNA extraction kit (Invitrogen, St. Louis, US) and snap frozen. RNA was extracted from the cell pellets using the MirVana RNA extraction kit (Invitrogen, St. Louis, US) as per manufacturer’s protocol. Genomic DNA removal was carried out using the DNA free kit (Thermo Fisher Scientific, Massachusetts, US) as per the manufacturer’s protocol. Two hundred ng/uL RNA (bESCs and hEECs) or 50 ng/uL RNA (bEECs) were reverse transcribed into cDNA using the High-Capacity cDNA Reverse Transcription Kit (Applied Biosystems, Massachusetts, US) as per the manufacturers protocol. Primers (IDT, Iowa, US) targeting bovine genes associated with early pregnancy gene expression changes in bovine endometrium and human genes associated with PDI (as determined by STRING DB analysis) were selected, including normaliser genes (bovine primers Table 1, human primers Table 2). Primers were designed using primer blast (28). qRT-PCR was carried out on a Roche Lightcycler 480 II (Roche, Basel, Switzerland) as per the standard Roche protocol with an annealing temperature of 60°C and ANOVA analysis of the 2^-ΔCt^ values were used to determine significant differences between the control and treated samples. ANOVA analysis and 2^-ΔΔCt^ graphs were created in GraphPad Prism software.

**Table 1.**
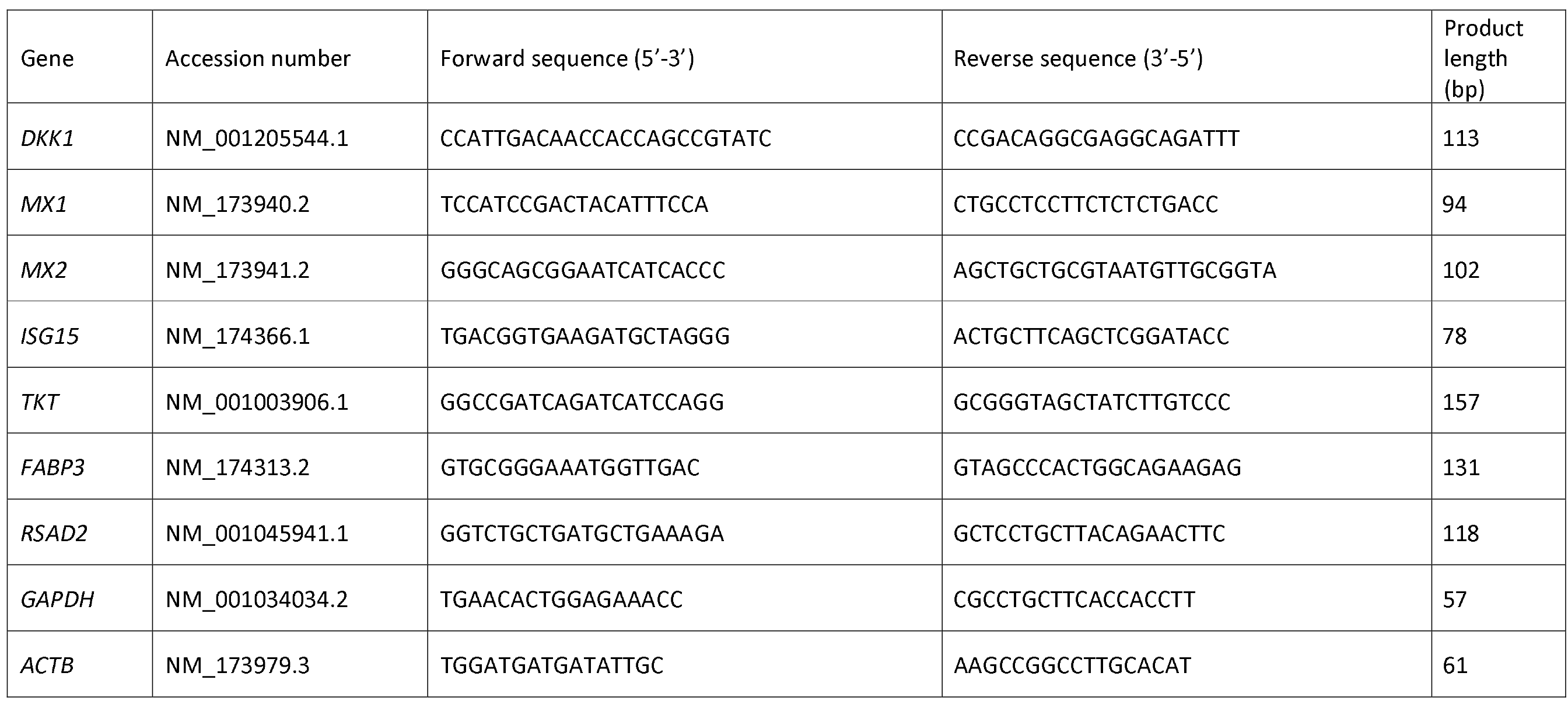
qRT-PCR primers designed to target bovine gene sequences specific to endometrial receptivity. Primers were designed using Primer Blast software (https://www.ncbi.nlm.nih.gov/tools/primer-blast/).

**Table 2.**
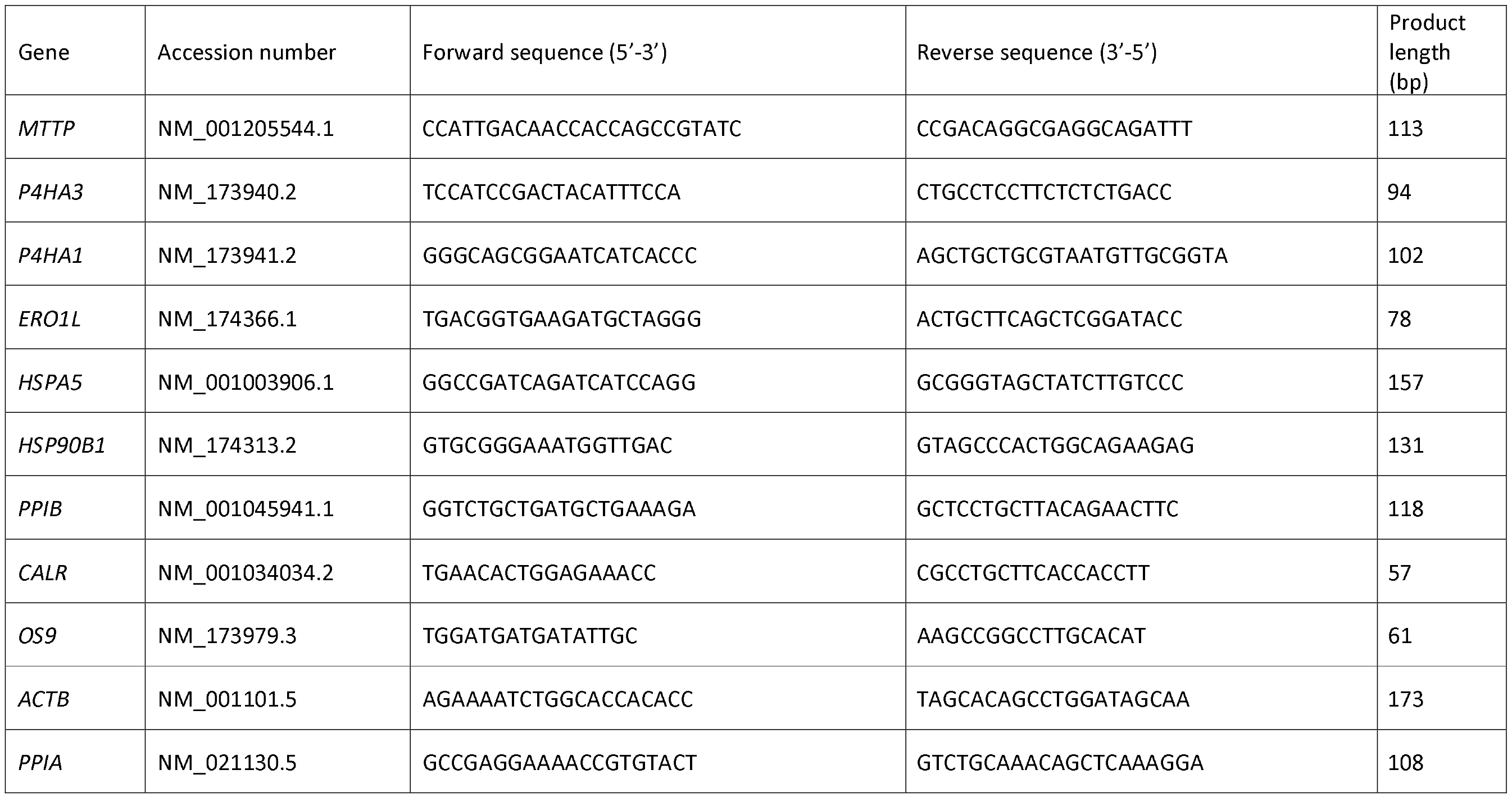
qRT-PCR primers designed to target human gene sequences specific to PDI. Primers were designed using Primer Blast software (https://www.ncbi.nlm.nih.gov/tools/primer-blast/).

### RNA sequencing and data analysis

Based on preliminary qRT-PCR data, the following samples were selected for RNA sequencing data analysis. bEEC and bESC (n=3) samples: 1) Control, 2) Vehicle control, 3) 1000 ng/ml rbPDI, and 4) 100 ng/ml roIFNT + 1000 ng/ml rbPDI. hEECs (n=3) samples: 1) Control 2) Vehicle control, 3) 1000 ng/ml rbPDI. RNA libraries were sequenced at the NGS Facility at the University of Leeds where they were prepared and sequenced as per their standard protocols using the Illumina TruSeq Stranded Total kit according to manufacturer’s guidelines, as described in detail previously (22). The RNA libraries were sequenced using the Illumina NextSeq 500 machine (Illumina, California, USA) with a single end 75 bp length read.

RNA sequencing data processing was conducted as described in detail in (22). Briefly, statistical test for differential gene expression was conducted via DESeq2 (29) with the cut-offs such as log2FoldChange >1 (or <-1) and padj <0.05. For principal component analysis (PCA) plotting of each group of samples, both protein-coding genes and long non-coding RNAs (lncRNAs) with RPKM value ≥ 1 in at least one sample were used and subsequently log2(RPKM+1) transformation and a quantile normalisation were applied. Overrepresentation enrichment analysis of differentially expressed protein-coding gene sets was executed using WebGestalt (30) for gene ontology terms and KEGG pathways. For gene ontology terms, biological process non-redundant datasets were chosen as functional database, and for both types of analyses, significance level was determined by FDR <0.05. Venn diagram analysis was performed using Venny (31).

### Implantation assays

#### Transfection validation

Following optimisation of transfection efficiency (Supplementary figure S1), *MNS1* knockdown was validated by plating Ishikawa cells (ECACC 99040201, passage 31) at a density of 100,000 cells per well in 12 well plates in triplicate (n=3) in 1mL complete human medium. After 24 hr the medium was aspirated, 1mL PBS added to wash cells and aspirated, and replenished with 800μL antibiotic-free human medium. A siRNA solution was made up for the *MNS1* siRNA (Horizon Discovery, UK) and the non-targeting siRNA (Horizon Discovery, UK) (1.4μL 100μM siRNA + 350μL OptiMEM). A lipofectamine solution was also prepared (900μL OptiMEM [Gibco, Massachusetts, US] + 18μL Lipofectamine 2000 [Invitrogen, St. Louis, US]). Following a 20-minute incubation together, the following solutions were added to each corresponding well: 1) Control (200μL OptiMEM only), 2) Vehicle control (100μL OptiMEM + 100μL Lipofectamine solution), 3) Non-targeting siRNA (100μL Lipofectamine solution + 100μL siRNA solution), 4) *MNS1* siRNA (100μL Lipofectamine solution + 100μL siRNA solution). After 4 hours incubation (37°C/5% CO_2_), medium was aspirated from the wells and replaced with complete human medium. Twenty-four hours after transfection, medium was aspirated, 1mL PBS added and aspirated to wash, and 700μL Qiazol (Qiagen, Germany) added to lyse cells. Cell lysate was collected in a 1.5mL Eppendorf, snap-frozen, and stored at -80°C until RNA extraction. RNA was extracted using the Qiagen miRNeasy Mini kit (Qiagen, Germany) with on-column DNA digestion (RNase-Free DNase set [Qiagen, Germany]) as per the manufacturers protocol and eluted in 50μL ultra-pure distilled water (Invitrogen, St. Louis, US). Purity and concentration of RNA was analysed using a NanoDrop ND-1000 Spectrophotometer (LabTech International, UK). RNA concentration was diluted to 200 ng/mL in ultra-pure distilled water (Invitrogen, St. Louis, US). Ten μL RNA was reverse transcribed into cDNA using the high-capacity cDNA synthesis kit (Applied Biosystems, Massachusetts, US) as described above. Two μL cDNA was plated into a 384 well clear plates (Bio-Rab Laboratories, California, US), in duplicate, per sample. Eight μL master mix, containing 0.25μL forward and reverse primers (IDT, Iowa, US), 2.5μ ultra-pure distilled water (Invitrogen, St. Louis, US), and 5μL SYBR Green (Roche, Switzerland) per well, was added. The plate was sealed with a microseal PCR plate sealing film (Bio-Rab Laboratories, California, US) and centrifuged briefly. The plate was then run using the standard Roche Lightcycler 480 mRNA programme with 40 cycles. Ct values were calculated using the Roche Lightcycler 480 software. The 2^-ΔΔCt^ method using *ACTB* as a normaliser gene, comparing all samples to the non-targeting samples, was used to determine fold change difference in expression. % knockdown of *MNS1* was calculated by taking the geometric mean of the 2^-ΔΔCt^ result and using the following equation: % knockdown = 100 - (100 * geomean of 2^-ΔΔCt^ result).

#### Implantation attachment assay

Ishikawa cells (ECACC 99040201, passage 37, n=5) were plated in 24-well plates (50,000 cells/well) in complete human medium. After 24 hours incubation at 37°C /5% CO_2_, each well had the medium aspirated, 1ml PBS (37°C) was added to each well to wash, and aspirated. To each well 400uL of antibiotic-free human medium was then added. An siRNA solution was made up for each siRNA (1.4μL 100uM siRNA [Horizon Discovery, UK] + 350μL OptiMEM [Gibco, Massachusetts, US]) and a lipofectamine solution (3μL Lipofectamine 2000 [Invitrogen, St. Louis, US] + 1.8mL OptiMEM). Following a 20-minute incubation, the following treatments were added to each corresponding well:

1. Control (100μL OptiMEM only)
2. Vehicle control (50μL OptiMEM + 50μL lipofectamine solution)
3. *MNS1* siGenome siRNA (50μL siRNA solution + 50μL lipofectamine solution)
4. SiGenome non-targeting siRNA (50μL siRNA solution + 50μL lipofectamine solution)
5. PBS vehicle control (20μL PBS + 80μL antibiotic-free human medium)
6. PDI (1μg/mL final concentration) (20μL 25μg/mL PDI stock + 80μL antibiotic-free human medium)

Plates were then returned to the incubator (37°C/5% CO_2_) for 48 hours before the assay.

Twenty-four hours prior to the assay, a 6-well plate was coated with 1mL of a 10% PVP (Sigma-Aldrich, Missouri, USA) solution for 30 minutes at room temperature. BeWo cells (ATCC CCL-98, passage 23) were lifted from a flask using trypsin (0.025%), pelleted at 500xg for 5 minutes, re- suspended in DMEM/F12 (Gibco, Massachusetts, US), and re-pelleted at 500xg for 5 minutes to remove all serum. The pellet was re-suspended in 3mL DMEM/F12, cells counted on a haematocytometer, and diluted to 166,000 cells/mL in DMEM/F12. Three mL cell suspension was added to each well after removing PVP solution and washing with 1mL PBS. The plate was then placed on an orbital shaker at 60rpm for 4 hours in an incubator (37°C/5% CO_2_). Each well was then pipetted up and down 10 times and the plate placed back into the incubator overnight.

After a 12-hour overnight incubation, BeWo spheroids were harvested by pipetting spheroids though a 100μM strainer (Corning, New York, US) above a 40μM strainer (Corning, New York, US) into a 50mL falcon. The original BeWo culture dish was washed with DMEM/F12, and residual spheroids also passed through the strainers. The flow-through was discarded and the 40μM strainer inverted over a 50mL falcon and backwashed with 1mL DMEM/F12.

Ishikawa cells after 48 hours had medium aspirated, 500μL DMEM/F12 added to remove traces of serum, aspirated and 450μL DMEM/F12 added. Fifty μL BeWo spheroids were added to the well of Ishikawa cells containing 450μL medium. The plate was them whole-well imaged on an EVOS light microscope (Thermo Fisher Scientific, Massachusetts, US) and then placed into an incubator (37°C/5% CO_2_) for 30 minutes to allow for attachment to occur. After 30 minutes, media was carefully removed, 500μL 10% neutral-buffered formalin (Epridia, Michigan, US) added gently and incubated at room temperature for 20 minutes to fix. After 20 minutes, the formalin was removed and 500μL of PBS was added. The whole well was then imaged again using an EVOS microscope. Using QuPath software (32) each spheroid was counted in the before-incubation and the after- incubation images for each well. Spheroids in the dark well rim were not counted, and clumps of spheroids had their number estimated. % attachment (spheroids remaining following fixing compared to number added to well) was calculated, and a paired students t-test performed for each treatment group compared to the relevant vehicle control.

### 2D microfluidics

#### Human endometrial epithelial cells

On day 1 of the experiment, human endometrial Ishikawa cells (passage 20, ECACC 99040201, n=3) were seeded at a density of 1,000,000/mL into IbiFlow 0.4 channel slides (Ibidi, Germany) (approximately 60μL to fill channel) in complete human medium, with the inlet and outlet marked. Cells were left to adhere for 4 hours in the incubator (37°C/5% CO_2_) before 100μL complete human medium was added to the inlet and the device returned to the incubator. Medium was replenished daily until day 4.

On day 3 the treatments were prepared in triplicate and put into 5mL leur lock syringes (Terumo, Japan) and placed into the incubator (37°C/5% CO_2_) overnight until attaching to devices as described below on day 4. Vehicle control treatment was 4.95mL complete human medium with 10% exosome depleted FBS + 50μL PBS. rbPDI treatment was 4.95mL complete human medium with 10% exosome depleted FBS + 50μL rbPDI (100 μg/mL).

On day 4, medium was removed from the outlet, and washed three times with 100μL PBS (37°C) added to the inlet and removed gently from the outlet. Complete human medium with 10% exosome depleted FBS (Gibco, Massachusetts, US) was replenished and device placed into incubator. The syringes were removed from the incubator, flicked to remove all bubbles, and placed securely into a NE-1600 syringe pump (New Era Pump Systems, New York, US) set up on an incubator shelf in a laminar flow hood. Each syringe had an Ibidi luer lock connector female (Ibidi, Germany) attached with approximately 10cm 0.8mm ID sterile silicone tubing (Ibidi, Germany). To the other end of the tubing an Ibidi elbow leur connector (Ibidi, Germany) was attached and media gently pushed through until no visible bubbles remained and a droplet of medium was present at the outlet of the elbow leur connector.

The device containing the cells was removed from the incubator and PBS added to the inlet until full. The droplet on the elbow connector was then connected to the full droplet of PBS in the device inlet and pressed down firmly at a 90-degree angle, then straightened to seal tightly, and repeated for all channels and syringes, as seen in Figures 2 and 3.

**Figure 1.**
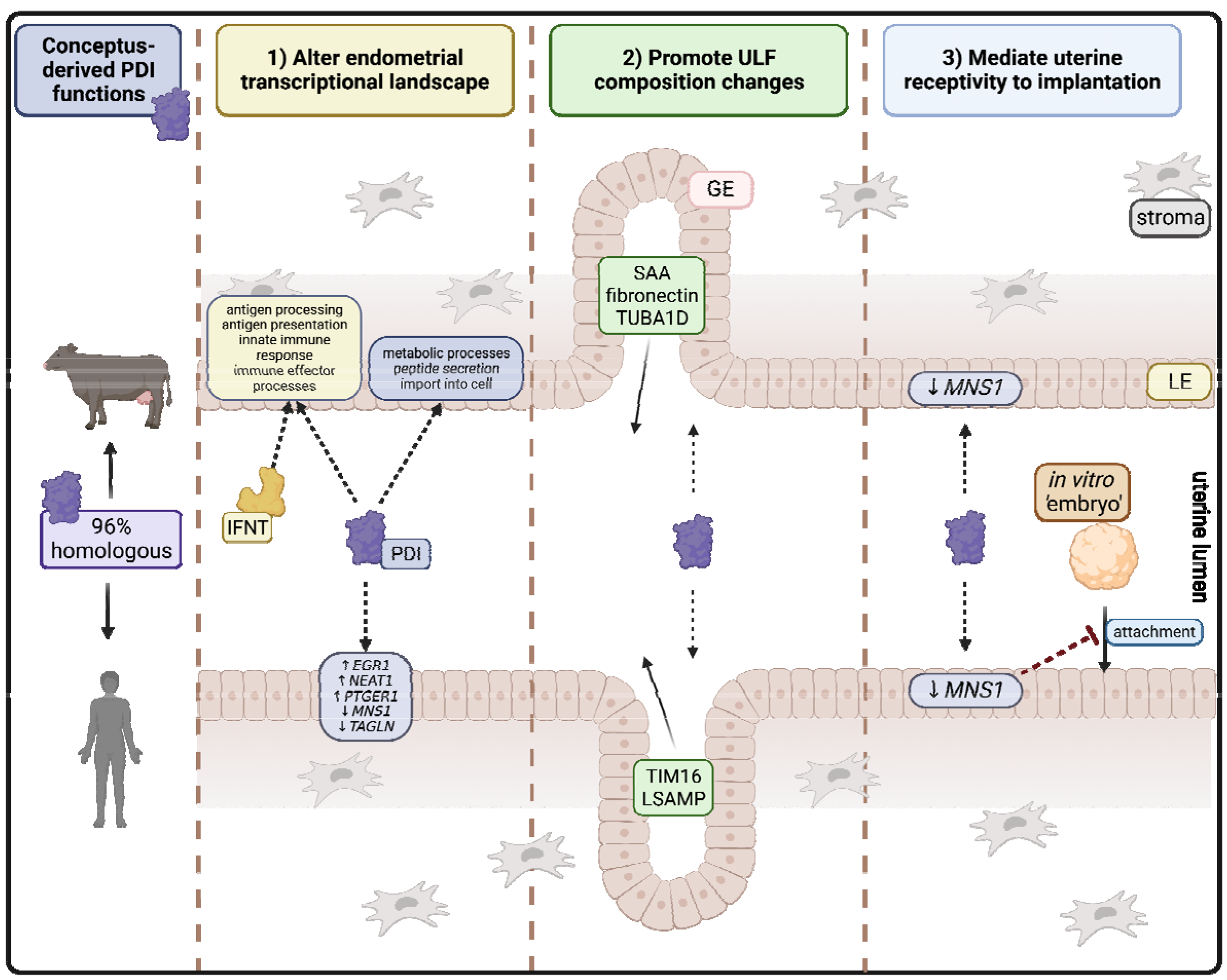
Graphical abstract summary of proposed actions of conceptus-derived PDI upon endometrial function. 1) PDI alters the endometrial landscape to support the actions of IFNT in the bovine endometrium and also alter specific pathways not induced by IFNT, and also induce a limited response in the human endometrium. 2) PDI promotes ULF composition changes to support conceptus development and nourishment pre-implantation. 3) PDI mediates uterine receptivity to implantation by reducing expression of MNS1 which may reduce attachment of the embryo to the endometrium at the incorrect time/location. LE = luminal epithelium, GE = glandular epithelial, IFNT = interferon tau, PDI = protein disulphide isomerase. Selected transcripts/GO terms and secreted proteins shown, full detail in supplementary tables. Figure created in BioRender.com.

**Figure 2.**
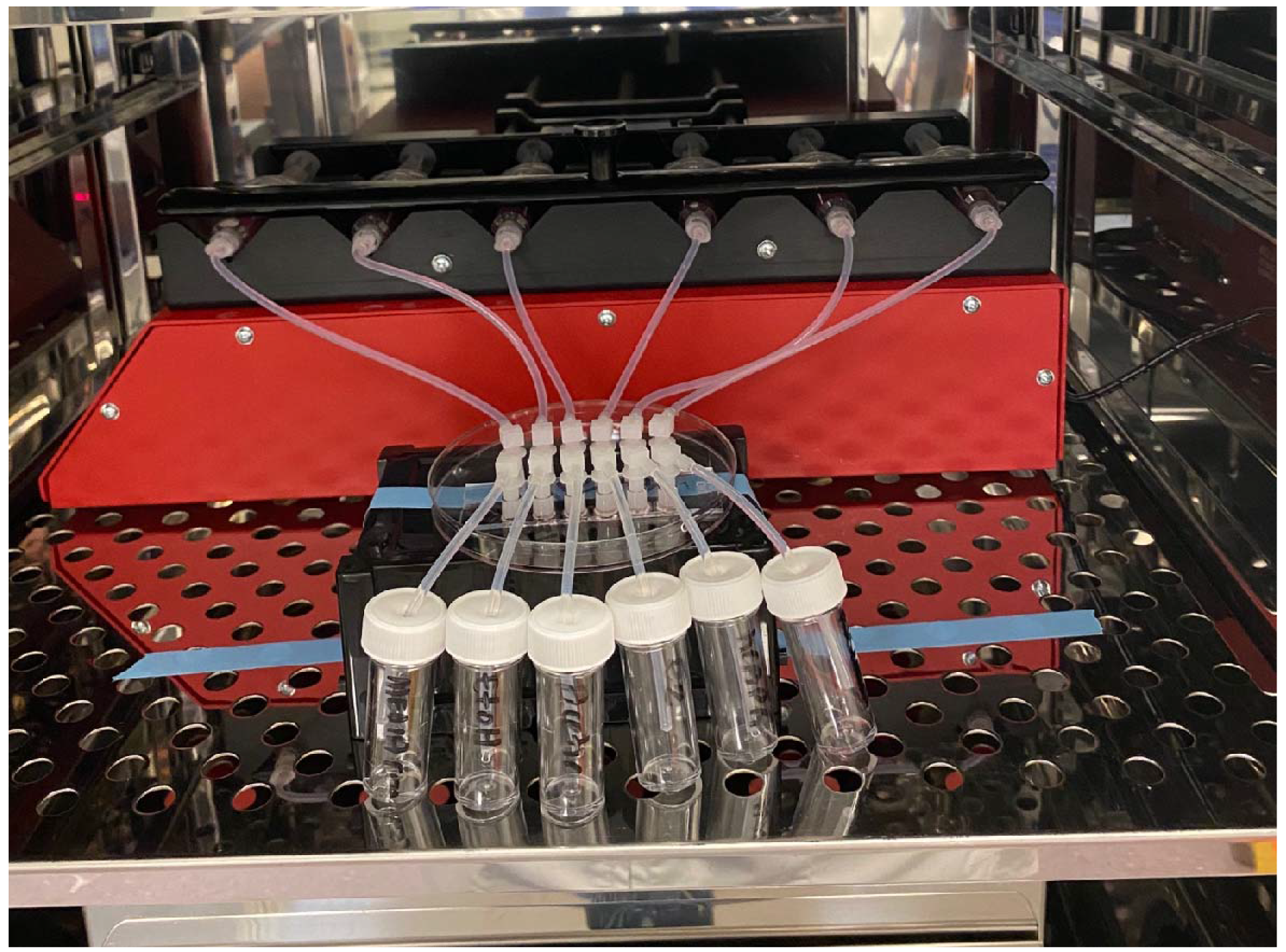
Representative example of 6-channel Ibidi device attached to syringe pump in incubator.

**Figure 3.**
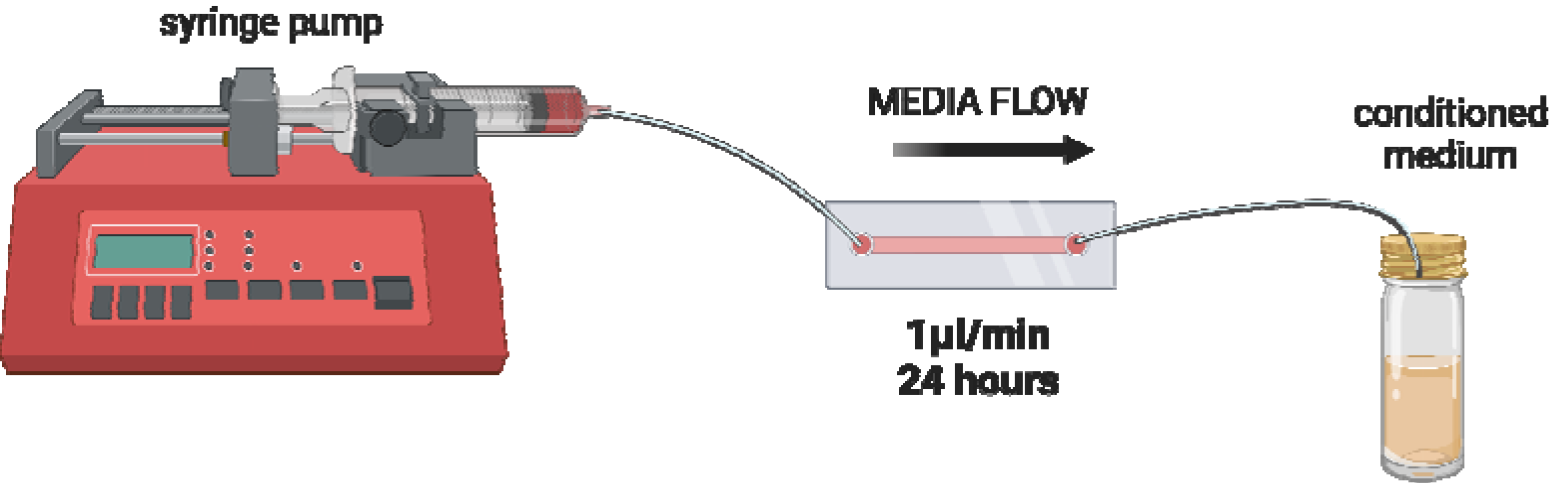
Graphical representation of microfluidics experimental setup. Syringe pump holds syringe filled with medium containing treatment is connected via tubing and connectors to a microfluidic device to flow medium through the channel (set flow rate 1μL/min).

Six 7mL bijous were prepared by pushing a hole in the lid. A second piece of 10cm sterile silicone 0.8mm ID tubing was connected to an elbow leur connector. A 1mL syringe (BD Plastipak, Michigan, US) with blunt needle (Sol-Millennium, Illionois, US) was used to fill the 10cm piece of tubing with PBS until a droplet of PBS emerged from the elbow leur connector. With the syringe still attached, the droplet of PBS on the elbow leur connector was joined with the droplet of PBS filling the outlet of the device channels and repeated for all six channels. The elbow leur connector was pressed firmly down into the outlet of the device at a 90-degree angle, whilst simultaneously removing the blunt needle. The elbow leur connector was then straightened to tightly connect with a firm seal. The end of the tubing was then placed through the hole into the 7mL bijou to collect the conditioned medium. The whole system was then taped securely with masking tape, and the shelf returned to the incubator (37°C/5% CO_2_). The system used is shown in Figures 2 and 3.

The syringe pump was switched on, set to a syringe internal diameter of 13mm, and a flow rate of 1uL/min, and set to run for 24 hours. This flow rate was selected because uterine tubal secretions was found to be 1.43mL/24 hours during dioestrus and 1.54mL/24 hours during oestrus in cattle (33), with the rate of 1uL/min equating to 1.44mL/24 hours. After 24 hours, the whole shelf was removed and placed into the laminar flow hood. The conditioned medium was collected from the 7mL bijous and transferred to 2mL sterile Eppendorf tubes and placed into the fridge until they could be processed. The Ibidi luer lock connector female was uncoupled from the syringes gently and the elbow connectors uncoupled from the devices. The device was then flushed through with PBS three times and then 0.025% trypsin added and placed into the incubator for 3 minutes. After three minutes the device was gently tapped to dislodge the cells and PBS added to the inlet to push the cells to the outlet where they were collected and transferred to a 1.5mL sterile Eppendorf. Cells were pelleted at 500xg for 5 minutes, re-suspended in 1mL PBS, and pelleted at 500xg for 5 minutes. PBS was aspirated and the pellet snap-frozen in liquid nitrogen before transferring to -80°C freezer.

#### Bovine endometrial epithelial cells

bEECs (n=3) were isolated as described above and seeded into devices after 27 days in culture at a density of 500,000 cells/mL in approximately 60μL complete bovine medium per channel. Cells were allowed to adhere for 4 hours in the incubator (37°C/5% CO_2_) and then 100μL complete bovine medium added to the inlet. Medium was replenished daily until day 4. On day 3 the treatments were prepared in triplicate and put into 5mL leur-lock syringes (Terumo, Japan) and placed into the incubator (37°C/5% CO_2_) overnight until attaching to devices. Vehicle control treatment was 4.95mL complete bovine medium with 10% charcoal stripped exosome depleted FBS + 50μL PBS. rbPDI treatment was 4.95mL complete bovine medium with 10% charcoal stripped exosome depleted FBS + 50μL rbPDI (100 μg/mL). On day 4, medium was removed from the outlet and the channel washed three times with 100μL PBS (37°C) added to the inlet and removed gently from the outlet. Complete bovine medium with 10% charcoal stripped exosome depleted FBS (Gibco, Massachusetts, US) was replenished and device placed into incubator. The devices were then attached to the syringes containing treatment in the pump system, and the experiment carried out under the same conditions and samples collected as described in for human microfluidic samples.

### Conditioned medium processing and proteomics analysis

The conditioned medium for both the human and bovine samples were processed from the fridge after collecting the cell pellets by centrifuging at 500xg for 10 minutes to remove cells, 2000xg for 10 minutes to remove cell debris, and microvesicles (MVs) pelleted at 14,500xg 4°C for 30 minutes. The supernatant conditioned medium (containing exosomes) was snap-frozen in liquid nitrogen and transferred to the -80°C freezer.

Following processing, the conditioned medium from both the human and bovine microfluidics were sent to Bristol Proteomics Facility for mass spectrometry analysis. Samples were depleted of bovine albumin using an albumin depletion kit, according to the manufacturer’s protocol (Thermo Fisher Scientific, UK). An equal volume of each depleted sample (equivalent to 30-40µg protein for the human samples, equivalent to 40-50µg protein for the bovine samples) was then digested with trypsin (1.25µg trypsin; 37°C, overnight), labelled with Tandem Mass Tag (TMT) ten plex reagents according to the manufacturer’s protocol (Thermo Fisher Scientific, UK) and the labelled samples pooled.

Pooled samples were desalted using a SepPak cartridge according to the manufacturer’s instructions (Waters, Massachusetts, USA). Eluate from the SepPak cartridge was evaporated to dryness and resuspended in buffer A (20 mM ammonium hydroxide, pH 10) prior to fractionation by high pH reversed-phase chromatography using an Ultimate 3000 liquid chromatography system (Thermo Fisher Scientific, UK). In brief, the sample was loaded onto an XBridge BEH C18 Column (130Å, 3.5 µm, 2.1 mm X 150 mm, Waters, UK) in buffer A and peptides eluted with an increasing gradient of buffer B (20 mM Ammonium Hydroxide in acetonitrile, pH 10) from 0-95% over 60 minutes. The resulting fractions (15 in total) were evaporated to dryness and resuspended in 1% formic acid prior to analysis by nano-LC MSMS using an Orbitrap Fusion Lumos mass spectrometer (Thermo Scientific).

High pH RP fractions were further fractionated using an Ultimate 3000 nano-LC system in line with an Orbitrap Fusion Lumos mass spectrometer (Thermo Scientific). In brief, peptides in 1% (vol/vol) formic acid were injected onto an Acclaim PepMap C18 nano-trap column (Thermo Scientific). After washing with 0.5% (vol/vol) acetonitrile 0.1% (vol/vol) formic acid peptides were resolved on a 250 mm × 75 μm Acclaim PepMap C18 reverse phase analytical column (Thermo Scientific) over a 150 minutes organic gradient, using 7 gradient segments (1-6% solvent B over 1min., 6-15% B over 58min., 15-32%B over 58min., 32-40%B over 5min., 40-90%B over 1min., held at 90%B for 6 minutes and then reduced to 1%B over 1min.) with a flow rate of 300 nl min^−1^. Solvent A was 0.1% formic acid and solvent B was aqueous 80% acetonitrile in 0.1% formic acid. Peptides were ionized by nano-electrospray ionization at 2.0kV using a stainless-steel emitter with an internal diameter of 30 μm (Thermo Scientific) and a capillary temperature of 300°C.

All spectra were acquired using an Orbitrap Fusion Lumos mass spectrometer controlled by Xcalibur 3.0 software (Thermo Scientific) and operated in data-dependent acquisition mode using an SPS- MS3 workflow. FTMS1 spectra were collected at a resolution of 120 000, with an automatic gain control (AGC) target of 200 000 and a max injection time of 50ms. Precursors were filtered with an intensity threshold of 5000, according to charge state (to include charge states 2-7) and with monoisotopic peak determination set to Peptide. Previously interrogated precursors were excluded using a dynamic window (60s +/-10ppm). The MS2 precursors were isolated with a quadrupole isolation window of 0.7m/z. ITMS2 spectra were collected with an AGC target of 10 000, max injection time of 70ms and CID collision energy of 35%.

For FTMS3 analysis, the Orbitrap was operated at 50 000 resolution with an AGC target of 50 000 and a max injection time of 105ms. Precursors were fragmented by high energy collision dissociation (HCD) at a normalised collision energy of 60% to ensure maximal TMT reporter ion yield. Synchronous Precursor Selection (SPS) was enabled to include up to 10 MS2 fragment ions in the FTMS3 scan.

The raw data files were processed and quantified using Proteome Discoverer software v2.1 (Thermo Scientific) and searched against the UniProt Bos taurus database (downloaded September 2020: 46224 entries) and the Uniprot Homo sapiens database (downloaded January 2021: 169297 entries) using the SEQUEST HT algorithm for the human samples. The raw data files were processed and quantified using Proteome Discoverer software v2.1 (Thermo Scientific) and searched against the UniProt Bos taurus database (downloaded October 2021: 37512 entries) and the Uniprot Homo sapiens database (downloaded January 2021: 169297 entries using the SEQUEST HT algorithm for the bovine samples. Peptide precursor mass tolerance was set at 10ppm, and MS/MS tolerance was set at 0.6Da. Search criteria included oxidation of methionine (+15.995Da), acetylation of the protein N-terminus (+42.011Da) and Methionine loss plus acetylation of the protein N-terminus (- 89.03Da) as variable modifications and carbamidomethylation of cysteine (+57.021Da) and the addition of the TMT mass tag (+229.163Da) to peptide N-termini and lysine as fixed modifications. Searches were performed with full tryptic digestion and a maximum of 2 missed cleavages were allowed. The reverse database search option was enabled, and all data was filtered to satisfy false discovery rate (FDR) of 5%.

The resulting list of proteins for each sample were then analysed as described by Aguilan *et al* in Excel to determine the fold change in protein abundance between conditioned medium from rbPDI treated cells compared to vehicle control (PBS only) and the associated p-value (34). First, any proteins which were present in the list but had no abundance values for any sample tested were removed from the list. Next, the data was transformed to a normal distribution using a log_2_ calculation followed by normalisation of the data by scaling each value against the average of all proteins within each sample. Next, the correlation slope of the average column for all samples compared with that of each sample was calculated. Then the log_2_ for each protein abundance was divided by the correlation slope for the sample, to further normalise the data by slope. Following this 2-step normalisation, any blank cells were randomly assigned a value which fits within the distribution (called deterministic minimum imputation) between 0-0.3. From the assigned value 2.5X the standard deviation of all values from that sample was removed. Fold changes were then calculated by taking the average of the vehicle control samples from the average of the rbPDI treated conditioned medium samples. Therefore, those proteins with a positive fold change are more abundant in the rbPDI conditioned medium samples compared to vehicle control, and the proteins with a negative fold change value are less abundant. P-value significance was calculated by paired two-tailed t-test, all comparisons with a p-value <0.05 were considered significantly different.

## RESULTS

### PDI is a highly conserved protein across mammals

The PDI amino acid sequence is highly conserved across representative placental mammals (>80% amino acid sequence conservation), and 96% conserved between human and bovine (Table 3).

**Table 3.** PDI percent amino acid identity matrix. . The darker shade represents higher levels of PDI amino acid sequence similarity, and lighter shade indicates lower similarity between species.

|  |  | Human | Mouse Lemur | Bushbaby | Mouse | Sheep | Cow | Dolphin | Dog | Horse | Microbat | Elephant | Armadillo | Wallaby | Opossum | Platypus | Chicken | Zebrafish |
| --- | --- | --- | --- | --- | --- | --- | --- | --- | --- | --- | --- | --- | --- | --- | --- | --- | --- | --- |
| Human | ENSP00000327801 | 100 | 95 | 93 | 94 | 95 | 96 | 77 | 94 | 97 | 94 | 88 | 94 | 71 | 87 | 85 | 84 | 73 |
| Mouse Lemur | ENSMICP00000001042 | 95 | 100 | 94 | 93 | 93 | 94 | 75 | 92 | 95 | 94 | 88 | 93 | 71 | 87 | 85 | 84 | 74 |
| Bushbaby | ENSOGAP00000002121 | 93 | 94 | 100 | 92 | 92 | 92 | 74 | 91 | 93 | 92 | 86 | 91 | 71 | 87 | 83 | 83 | 72 |
| Mouse | ENSMUSP00000026122 | 94 | 93 | 92 | 100 | 94 | 94 | 74 | 92 | 96 | 93 | 87 | 93 | 71 | 86 | 84 | 84 | 73 |
| Sheep | ENSOARP00000019364 | 95 | 93 | 92 | 94 | 100 | 99 | 79 | 94 | 97 | 95 | 88 | 94 | 70 | 86 | 84 | 84 | 73 |
| Cow | ENSBTAP00000007943 | 96 | 94 | 92 | 94 | 99 | 100 | 80 | 94 | 98 | 96 | 88 | 94 | 71 | 87 | 85 | 84 | 72 |
| Dolphin | ENSTTRP00000011641 | 77 | 75 | 74 | 74 | 79 | 80 | 100 | 76 | 76 | 77 | 70 | 74 | 52 | 69 | 67 | 66 | 57 |
| Dog | ENS CAF P00000008777 | 94 | 92 | 91 | 92 | 94 | 94 | 76 | 100 | 96 | 95 | 86 | 92 | 71 | 85 | 83 | 84 | 73 |
| Horse | ENSECAP00000007958 | 97 | 95 | 93 | 96 | 97 | 98 | 76 | 96 | 100 | 97 | 94 | 94 | 72 | 91 | 86 | 87 | 75 |
| Microbat | ENSM LUP00000017209 | 94 | 94 | 92 | 93 | 95 | 96 | 77 | 95 | 97 | 100 | 88 | 93 | 70 | 86 | 84 | 85 | 73 |
| Elephant | ENSLAFP00000001175 | 88 | 88 | 86 | 87 | 88 | 88 | 70 | 86 | 94 | 88 | 100 | 90 | 67 | 83 | 81 | 83 | 69 |
| Armadillo | ENSDNOP00000009104 | 94 | 93 | 91 | 93 | 94 | 94 | 74 | 92 | 94 | 93 | 90 | 100 | 70 | 88 | 85 | 84 | 73 |
| Wallaby | ENSMEUP00000015288 | 71 | 71 | 71 | 71 | 70 | 71 | 52 | 71 | 72 | 70 | 67 | 70 | 100 | 78 | 70 | 66 | 58 |
| Opossum | ENSMODP00000003645 | 87 | 87 | 87 | 86 | 86 | 87 | 69 | 85 | 91 | 86 | 83 | 88 | 78 | 100 | 82 | 80 | 70 |
| Platypus | ENSOANP00000009770 | 85 | 85 | 83 | 84 | 84 | 85 | 67 | 83 | 86 | 84 | 81 | 85 | 70 | 82 | 100 | 79 | 71 |
| Chicken | ENSGALP00000056764 | 84 | 84 | 83 | 84 | 84 | 84 | 66 | 84 | 87 | 85 | 83 | 84 | 66 | 80 | 79 | 100 | 70 |
| Zebrafish | ENSDARP00000138657 | 73 | 74 | 72 | 73 | 73 | 72 | 57 | 73 | 75 | 73 | 69 | 73 | 58 | 70 | 71 | 70 | 100 |

### rbPDI alters early-pregnancy associated gene expression in bESCs

When treated with rbPDI at the highest concentration used (1000 ng/ml rbPDI), there was a significant increase (p<0.05) in the expression of transcripts *MX1*, *MX2*, *RSAD2*, and *ISG15* (Figure 4). As expected, the addition of roIFNT in combination with rbPDI also lead to the significant increase in the expression of *MX1*, *MX2*, *RSAD2*, and *ISG15* in bESCs at all concentrations tested, although at much higher fold changes of expression than rbPDI treatment alone (Figure 5). Neither rbPDI alone **Error! Reference source not found.**(Figure 4), or in combination with roIFNT (Figure 5) significantly altered the expression of the transcripts *FABP3* or *DKK1* at the concentrations (10/100/1000 ng/mL rbPDI with/without 100 ng/mL roIFNT) or duration of treatment (48 hours) tested.

**Figure 4.**
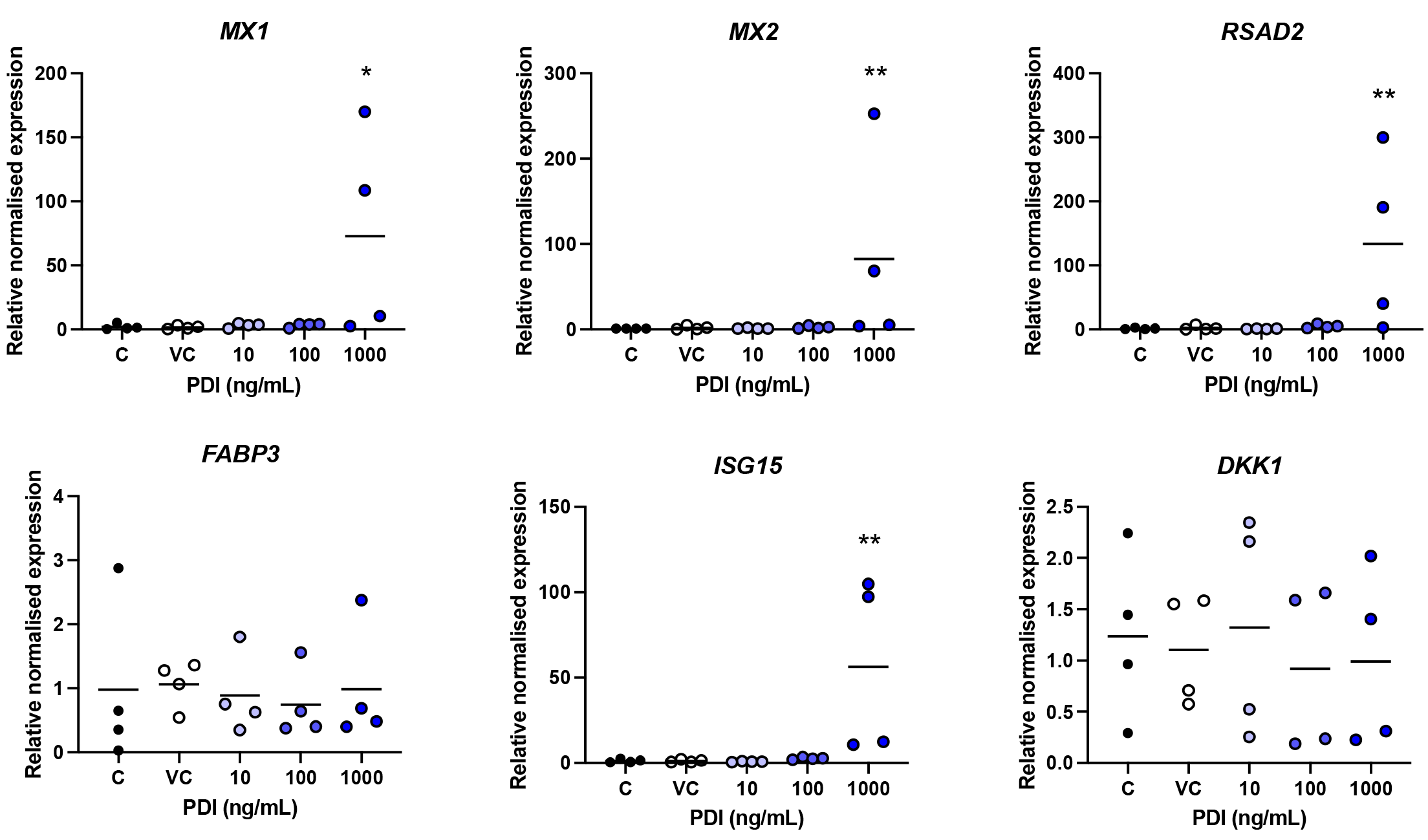
Relative expression of interferon-stimulated genes in bovine endometrial stromal cells following treatment with rbPDI (1000 ng/ml) for 24 hours (n=4). Relative gene expression to the vehicle control samples determined by qRT-PCR and the 2-DDCt method (1), with ACTB and GAPDH as normaliser genes. Significance determined by ANOVA analysis, * P<0.05, ** P<0.01. Figure created in Graphpad Prism. Line at mean.

**Figure 5.**
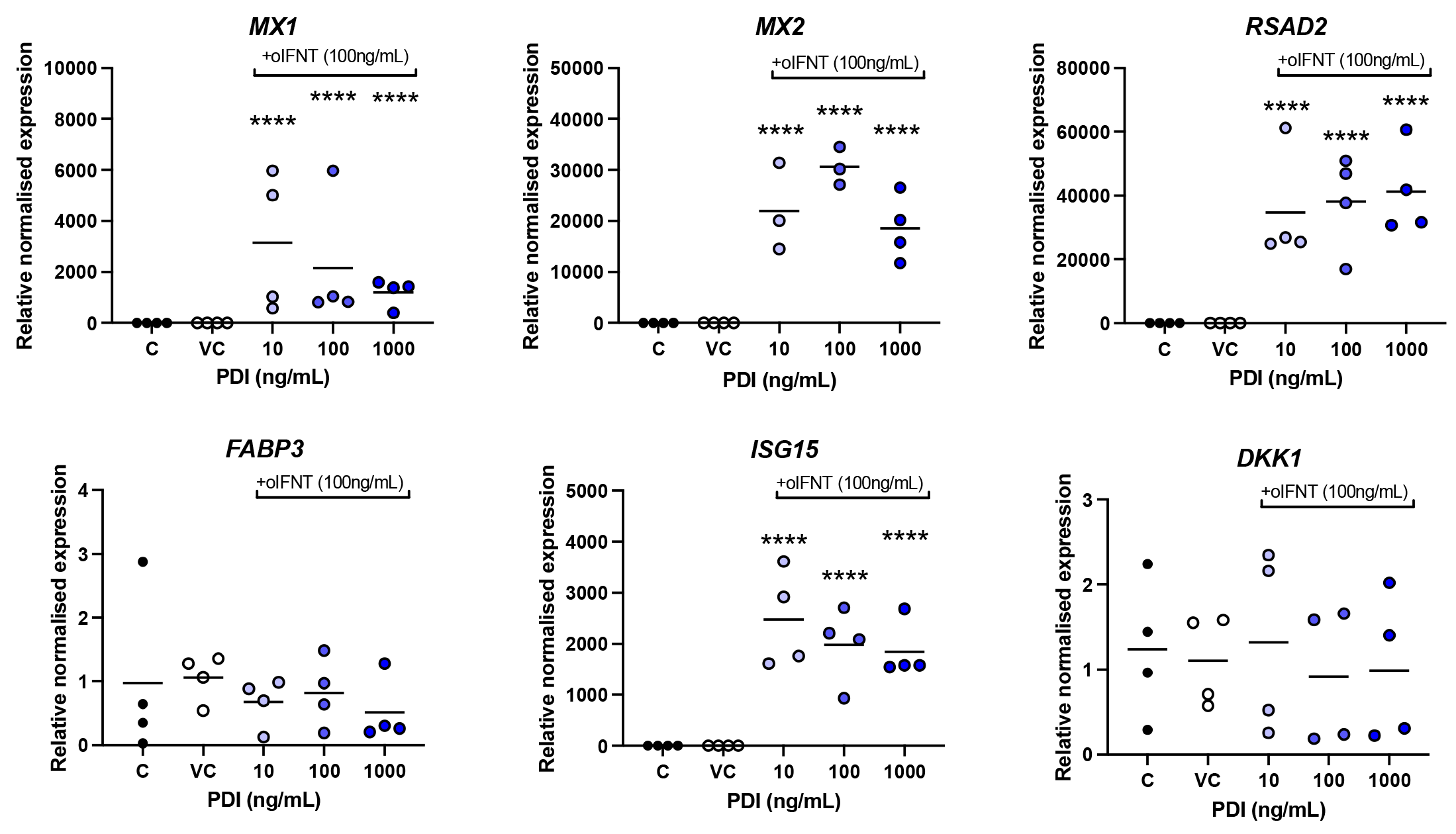
Relative expression of interferon-stimulated genes in bovine endometrial stromal cells following treatment with rbPDI (1000 ng/ml) in combination with roIFNT (100 ng/ml) for 24 hours (n=4). Relative gene expression to the vehicle control samples determined by qRT-PCR and the 2-DDCt method (1), with ACTB and GAPDH as normaliser genes. Significance determined by ANOVA analysis, **** P<0.0001. Figure created in Graphpad Prism. Line at mean.

### rbPDI does not alter expression of any PDI-associated transcripts in hEECs

As seen in Figure 6, rbPDI treatment of hEECs did not significantly alter the expression of any of the PDI-associated protein’s mRNA transcripts (*P4HA1, PPIB, CALR, HSPA5, OS9, P4HA3*) at the concentrations (10/100/1000 ng/mL) and treatment duration (48 hours) tested.

**Figure 6.**
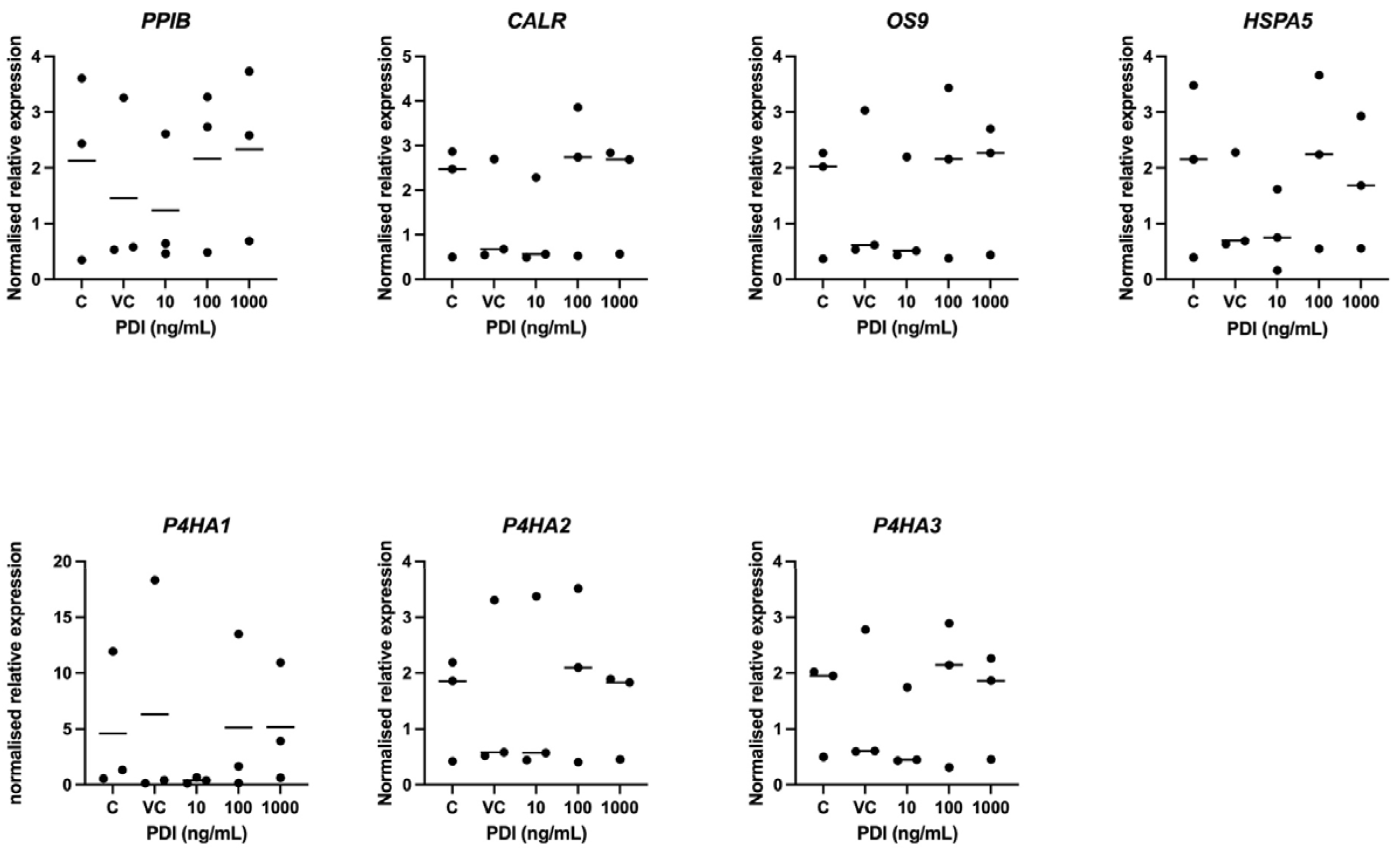
Relative expression of PDI-associated genes in human Ishikawa endometrial epithelial cells following treatment with r bPDI. rbPDI concentrations varied (10, 100, 1000ng/mL) and cells treated for 24 hours (n=3). Relative gene expression to the vehicle control samples determined by qRT-PCR and the 2^-ᡸᡸCt^ method (1), with ACTB and GAPDH as normaliser genes. Significance determined by ANOVA analysis, ns = not significant p>0.05. Figure created in Graphpad Prism. Line at mean.

### rbPDI induces a transcriptional response in bEEC and bESCs

Based on the qRT-PCR data demonstrating that the transcriptional response to rbPDI in bESCs occurred in response to the highest concentration of rbPDI tested (1000 ng/ml), the samples treated with 1000 ng/ml rbPDI were used for RNA sequencing. rbPDI induced a transcriptional response in both bESCs and bEECs distinct from controls, as seen in Figure 7. The PCA plots show that the control samples cluster towards the left, whereas the samples treated with rbPDI cluster more towards the right of the plot. rbPDI treatment altered the expression of 448 transcripts compared to VC in bEECs, including 4 upregulated lncRNAs (Supplementary table S1). Of the protein coding transcripts, 365 were upregulated and 83 downregulated compared to VC. Twenty-four GO terms (Figure 8, Supplementary table S2) and 38 KEGG pathways were enriched within the DEGs compared to VC (Figure 9, Supplementary table S3). In bESCs, rbPDI treatment altered the expression of 306 DEGs compared to VC, all of which were protein coding transcripts (Supplementary table S4). Three hundred protein coding transcripts were upregulated and 6 downregulated compared to VC.

**Figure 7.**
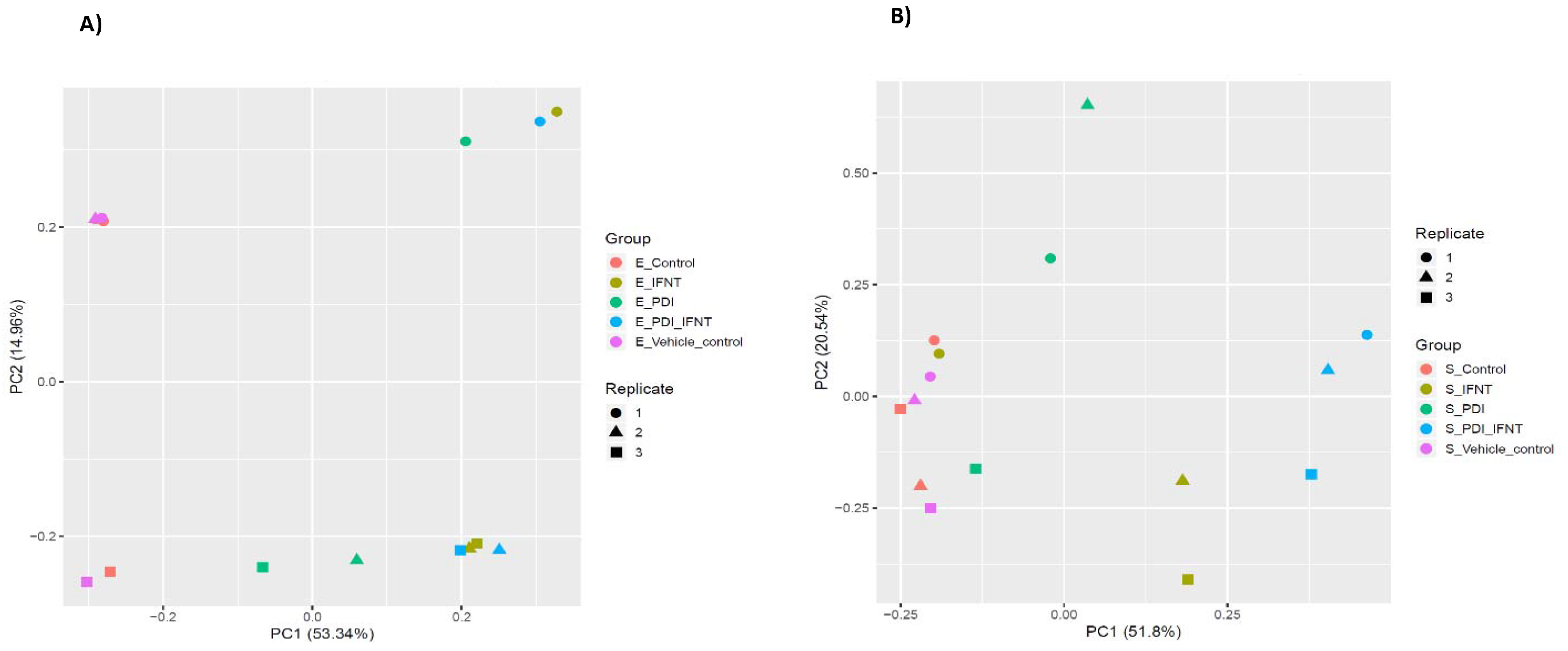
Principal component analysis describing the transcriptional effect of rbPDI and roIFT. Including rbPDI treatment alone, roIFNT treatment alone, and both in combination compared to control and vehicle control samples in A) bovine endometrial epithelial cells or B) bovine endometrial stromal cells (n=3). Data points indicate each replicate and data point colours indicate treatment groups. Control samples (_Control), vehicle control samples (_vehicle_control), roIFNT treatment (_IFNT), rbPDI treatment (_PDI), and rbPDI + roIFNT in combination (_PDI_IFNT).

**Figure 8.**
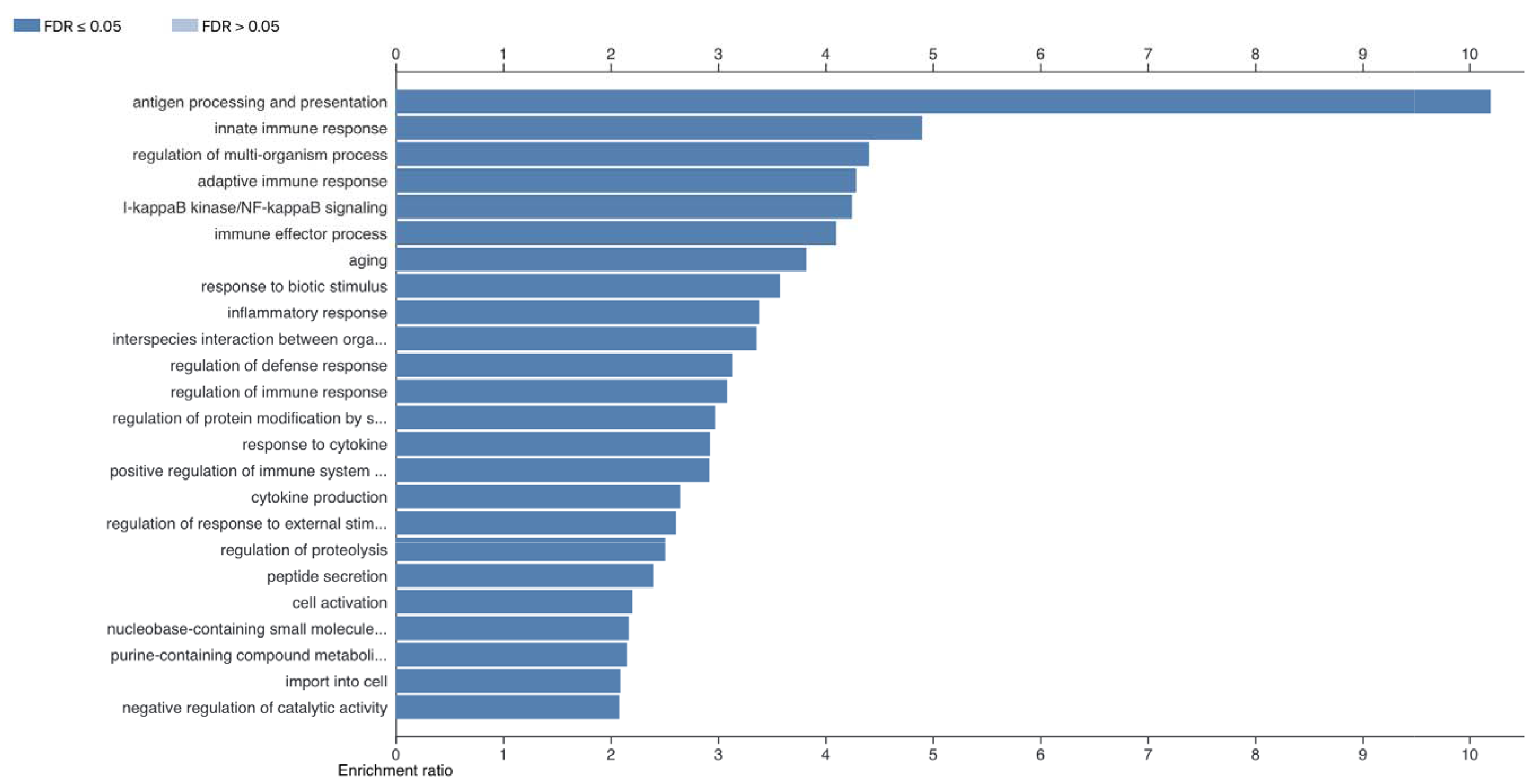
Enriched GO terms in DEGs specific to bEECs treated with rbPDI. DEGs including uncharacterised transcripts compared to VC samples (padj<0.05) analysed by WebGestalt to determine enriched go terms (FDR <0.05). Full data in Supplementary table 2.

**Figure 9.**
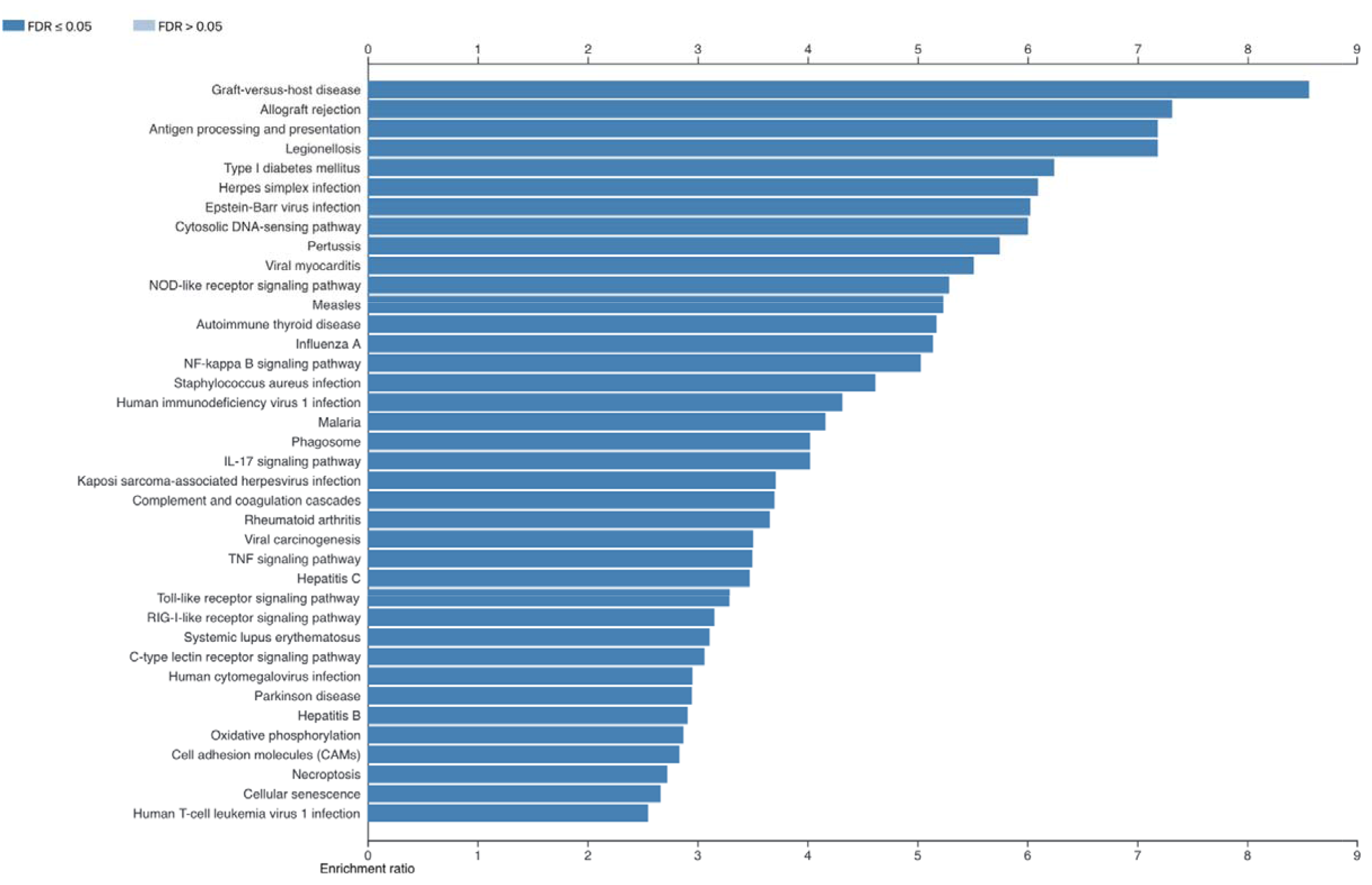
Enriched KEGG pathways in DEGs specific to bEECs treated with rbPDI. DEGs including uncharacterised transcripts compared to VC samples (padj<0.05) analysed by WebGestalt to determine enriched go terms (FDR <0.05). Full data in Supplementary table 3.

Twenty-two GO terms (Figure 10, Supplementary table S5) and 26 KEGG pathways were enriched with the DEGs compared to VC (Figure 11, Supplementary table S6). Venn diagram analysis demonstrated the transcriptional response induced by rbPDI broadly differed between bEECs and bESCs (Figure 12, Supplementary table S7), although 67 transcripts were commonly altered in both cell types. Downstream analysis on the 67 transcripts revealed 27 overrepresented GO terms (Figure 13, Supplementary table S8), and 14 enriched KEGG pathways (Figure 14, Supplementary table S9).

**Figure 10.**
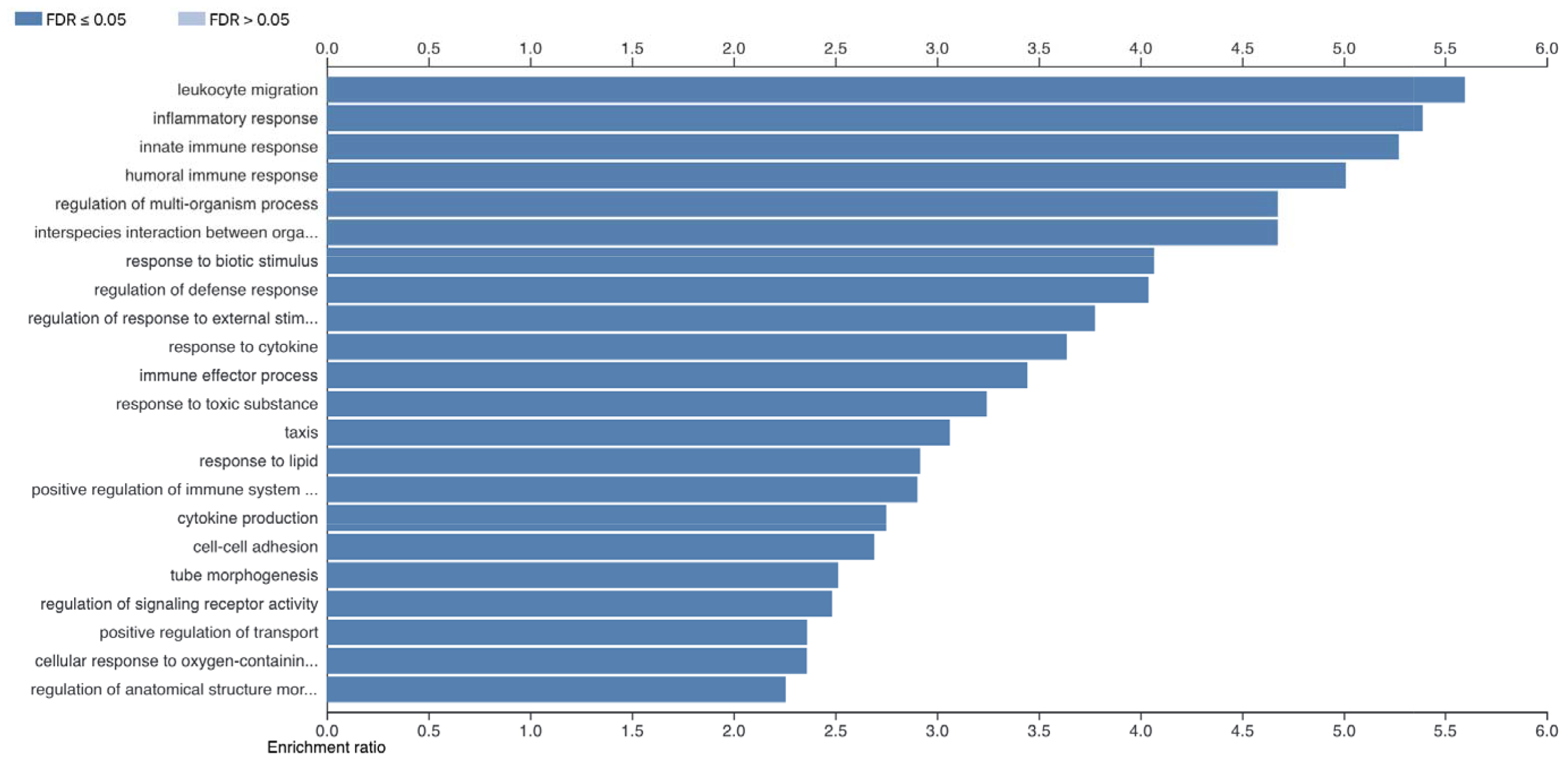
Enriched GO terms in DEGs specific to bESCs treated with rbPDI. DEGs including uncharacterised transcripts compared to VC samples (padj<0.05) analysed by WebGestalt to determine enriched go terms (FDR <0.05). Full data in Supplementary table 5.

**Figure 11.**
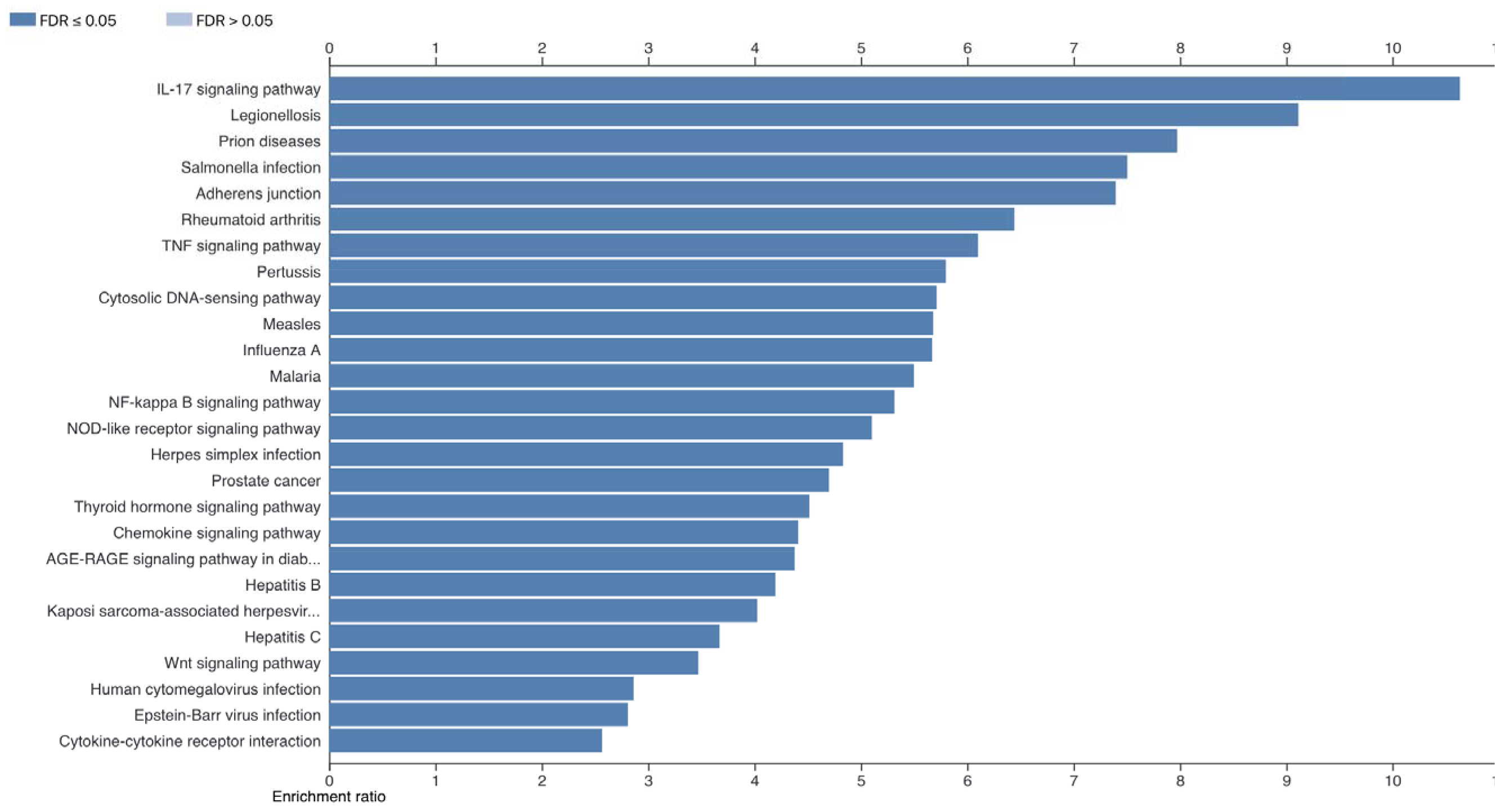
Enriched KEGG pathways in DEGs specific to bESCs treated with rbPDI. DEGs including uncharacterised transcripts compared to VC samples (padj<0.05) analysed by WebGestalt to determine enriched go terms (FDR <0.05). Full data in Supplementary table 6.

**Figure 12.**
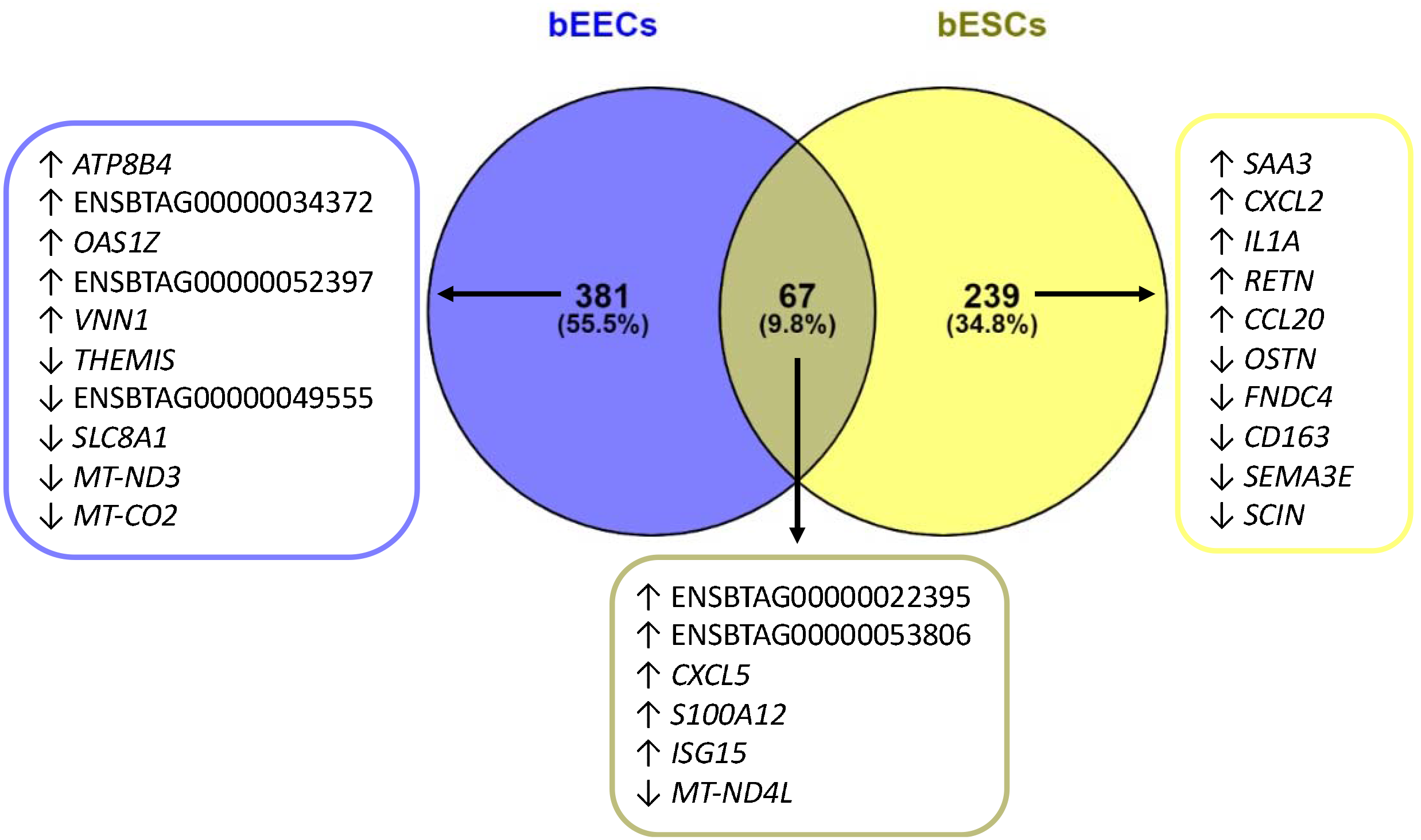
Venn diagram analysis of rbPDI treatment induced DEGs in bEECs vs bESCs. rbPDI compared to vehicle control samples (n=3) in bovine endometrial epithelial cells (bEECs) and stromal cells (bESCs). Top 5 up or downregulated transcripts in each group included. Full data Supplementary table 7.

**Figure 13.**
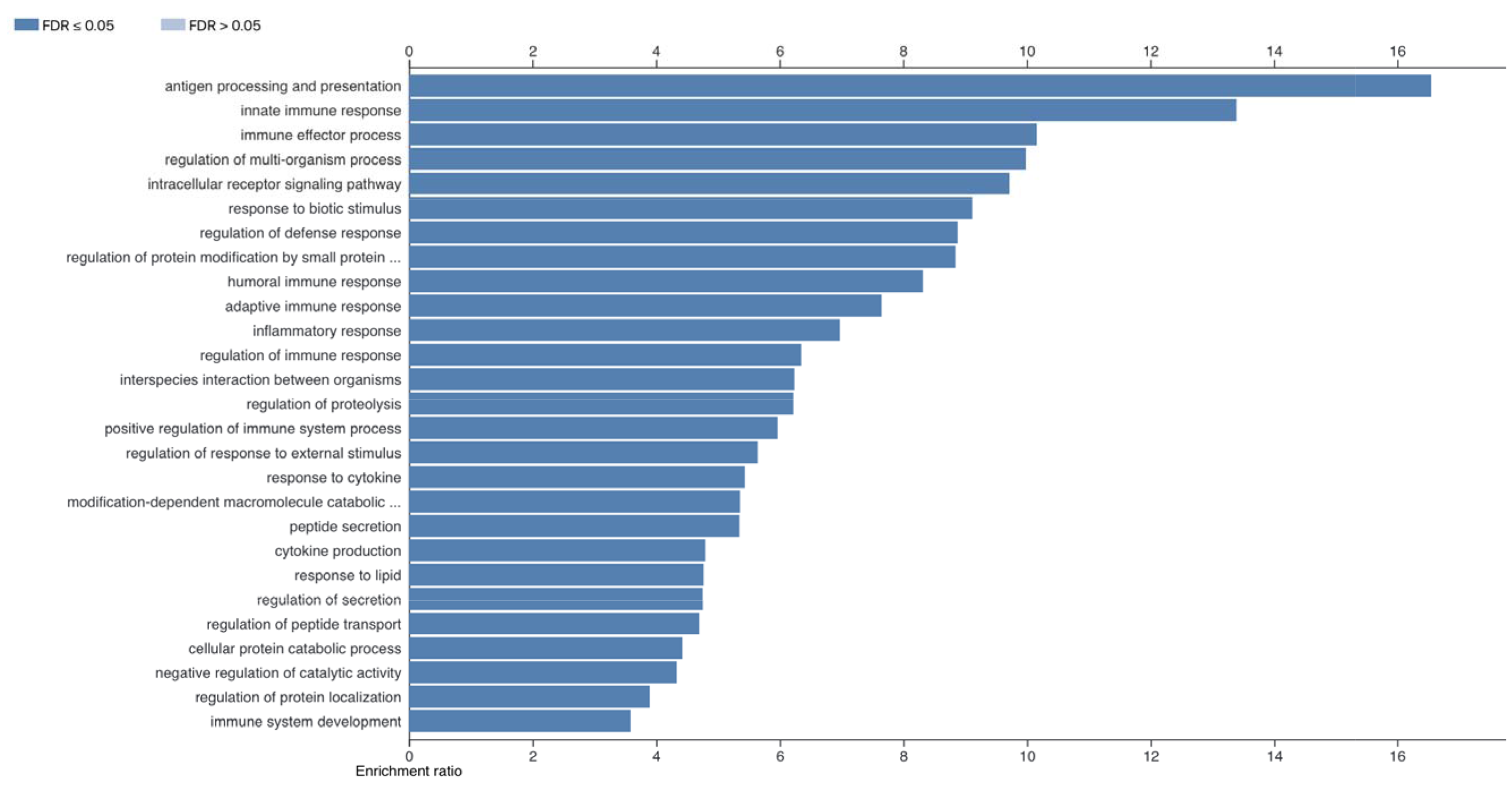
Enriched GO terms in 67 DEGs specific to both bEECs and bESCs treated with rbPDI. DEGs including uncharacterised transcripts compared to VC samples (padj<0.05) analysed by WebGestalt to determine enriched go terms (FDR <0.05). Full data in Supplementary table 8.

**Figure 14.**
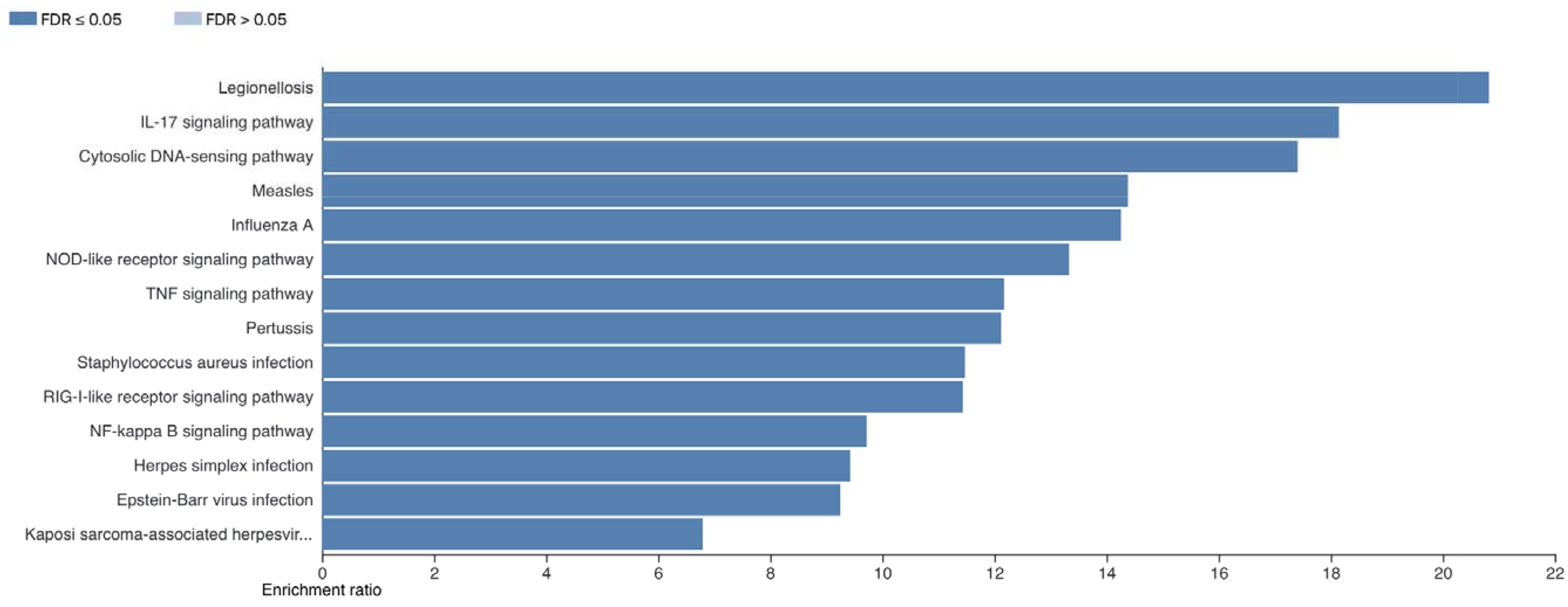
Enriched KEGG pathways in 67 DEGs specific to both bEECs and bESCs treated with rbPDI. DEGs including uncharacterised transcripts compared to VC samples (padj<0.05) analysed by WebGestalt to determine enriched go terms (FDR <0.05). Full data in Supplementary table 9.

### rbPDI alone, and in combination with roIFNT, induces a distinct transcriptional response to roIFNT alone in bESC and bEECs

bEECs treated with roIFNT in combination with rbPDI led to 1415 DEGs. Of the 1385 altered protein coding transcripts, 974 were upregulated and 411 downregulated compared to VC. Of the altered 30 lncRNAs, 23 were upregulated and 7 downregulated compared to VC (Supplementary table S10). Forty-three GO terms and 36 KEGG pathways were enriched among the DEGs compared to VC (Supplementary table S11 + S12).

Venn diagram analysis (Figure 15A, Supplementary table S13 + S19) determined that 341 transcripts were commonly differentially expressed in bEECs in response to all three treatments of: roIFNT only (22), rbPDI only, and rbPDI and roIFNT in combination when compared to VC samples. rbPDI and roIFNT in combination elicited the differential expression of 240 transcripts which were not altered in roIFNT or rbPDI only treatments. Eighty-nine transcripts were differentially expressed in response to PDI alone and in combination with roIFNT, but not by roIFNT alone. Additionally, 17 transcripts were uniquely differentially expressed in response to rbPDI, but not when treated in combination with roIFNT. Further analysis of the 341 DEGs commonly altered by both rbPDI and roIFNT alone and in combination in bEECs compared to VC revealed 13 enriched GO terms and 23 enriched KEGG pathways (Supplementary table S14 + S15). Further analysis of the 346 DEGs commonly altered by both rbPDI alone and rbPDI in combination with roIFNT, but not roIFNT alone in bEECs compared to VC revealed 21 enriched GO terms (Figure 16) and 11 enriched KEGG pathways (Figure 17). bESCs treated with rbPDI together with roIFNT elicited the differential expression of 2020 transcripts, including 32 lncRNAs and 1988 protein-coding transcripts. Of the lncRNAs, 20 were upregulated and 12 downregulated compared to VC, and of the protein-coding transcripts, 1367 were upregulated and 621 downregulated compared to VC (Supplementary table S16). Fifty-three GO terms and 62 KEGG pathways were enriched in the DEGs compared to VC (Supplementary table S17 + S18).

**Figure 15.**
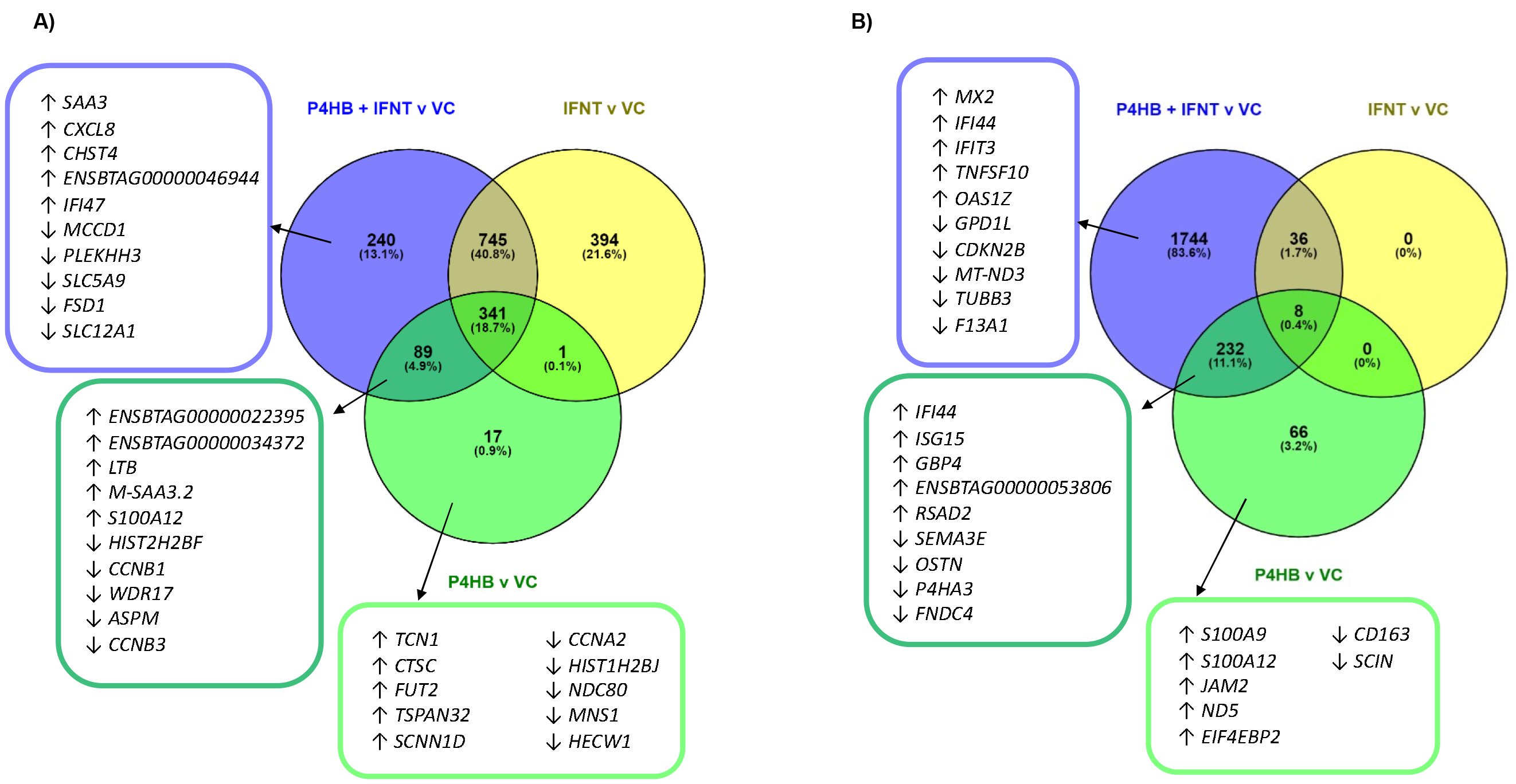
Venn diagram analysis of DEGs induced in bovine endometrial cells treated with roIFNT. (2)**, rbPDI, or rbPDI in combination with roIFNT** (2). A) bEECs or B) bESCs when compared to vehicle control samples. Full data in Supplementary table 13+19.

**Figure 16.**
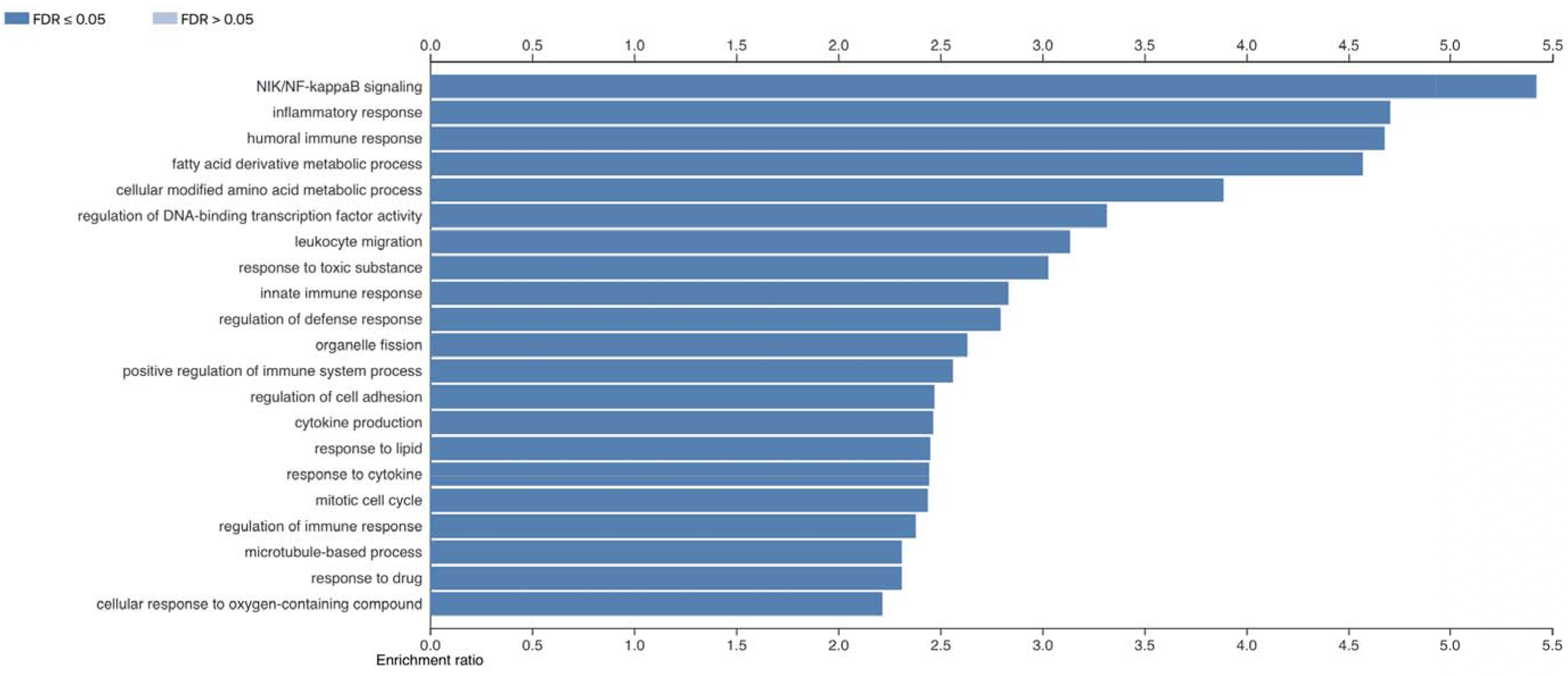
Enriched GO terms of DEGs specific to bEECs treated with rbPDI vs VC or rbPDI + roIFNT, but not induced by roIFNT, v s VC. DEGs including uncharacterised transcripts compared to VC samples (padj<0.05) analysed by WebGestalt to determine enriched go terms (FDR <0.05).

**Figure 17.**
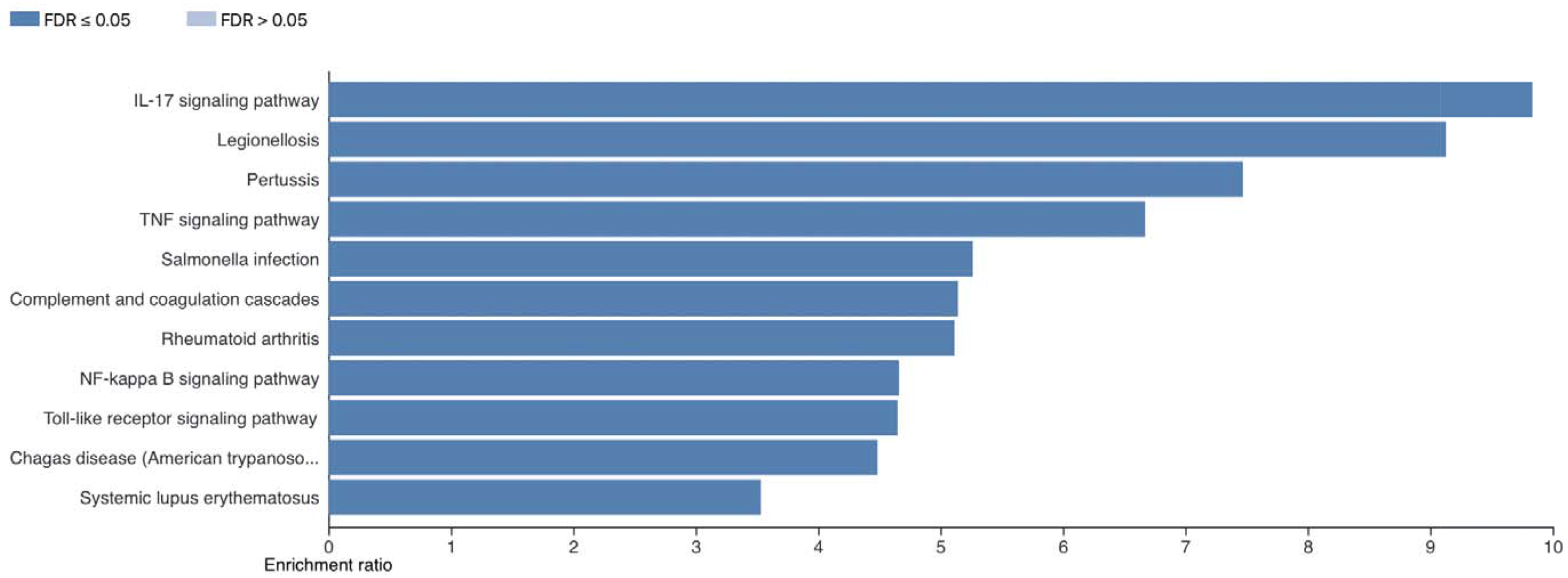
Enriched KEGG pathways of DEGs specific to bEECs treated with rbPDI vs VC or rbPDI + roIFNT, but not induced by roIF NT, vs VC. DEGs including uncharacterised transcripts compared to VC samples (padj<0.05) analysed by WebGestalt to determine enriched KEGG pathways (FDR <0.05).

In bESCs, a comparison of roIFNT and rbPDI treatment in bESCs revealed that of the 44 DEGs altered by roIFNT and the 306 DEGs altered by rbPDI compared to VC, 8 transcripts were commonly altered by both treatments alone and in combination (Figure 15B, Supplementary table S19). Venn analysis showed that the 44 transcripts altered by roIFNT treatment were also altered by roIFNT treatment in combination with rbPDI. Eight transcripts were commonly altered by all three treatment groups, and 232 of the transcripts altered by rbPDI alone were also altered when treated with rbPDI in combination with roIFNT. Sixty-six transcripts were uniquely altered by PDI treatment alone, but not in combination with roIFNT or roIFNT alone. Further analysis on the 8 DEGs altered in response to all three treatments did not result in any significantly enriched GO terms or KEGG pathways.

### hEECs have a limited transcriptional response to rbPDI

rbPDI elicited a transcriptional response in hEECs *in vitro* compared to vehicle control treatment, based on the PCA plot (Figure 18). The treated samples clustered towards the right of the PCA plot and the controls on the left. Forty-nine transcripts (7 of which are lncRNAs) were altered by rbPDI compared to VC (Supplementary table S20). Twenty-six of the protein-coding transcripts were upregulated and 16 downregulated compared to VC. No significantly enriched GO terms or KEGG pathways were identified. Very few transcripts were conserved in the transcriptional response to rbPDI between bovine and human compared to VC. However, there was a shared response to rbPDI in both bovine and human endometrial epithelial cells. A Venn comparison of DEGs altered by rbPDI treatment compared to VC samples in bEECs, bESCs, and hEECs demonstrated that of the 49 characterised transcripts altered in hEECs, 1 transcript (*MNS1)* was also altered in bEECs, and 2 transcripts (*NLGN2* and *ULK1*) were also altered in bESCs (Figure 19, Supplementary table S21).

**Figure 18.**
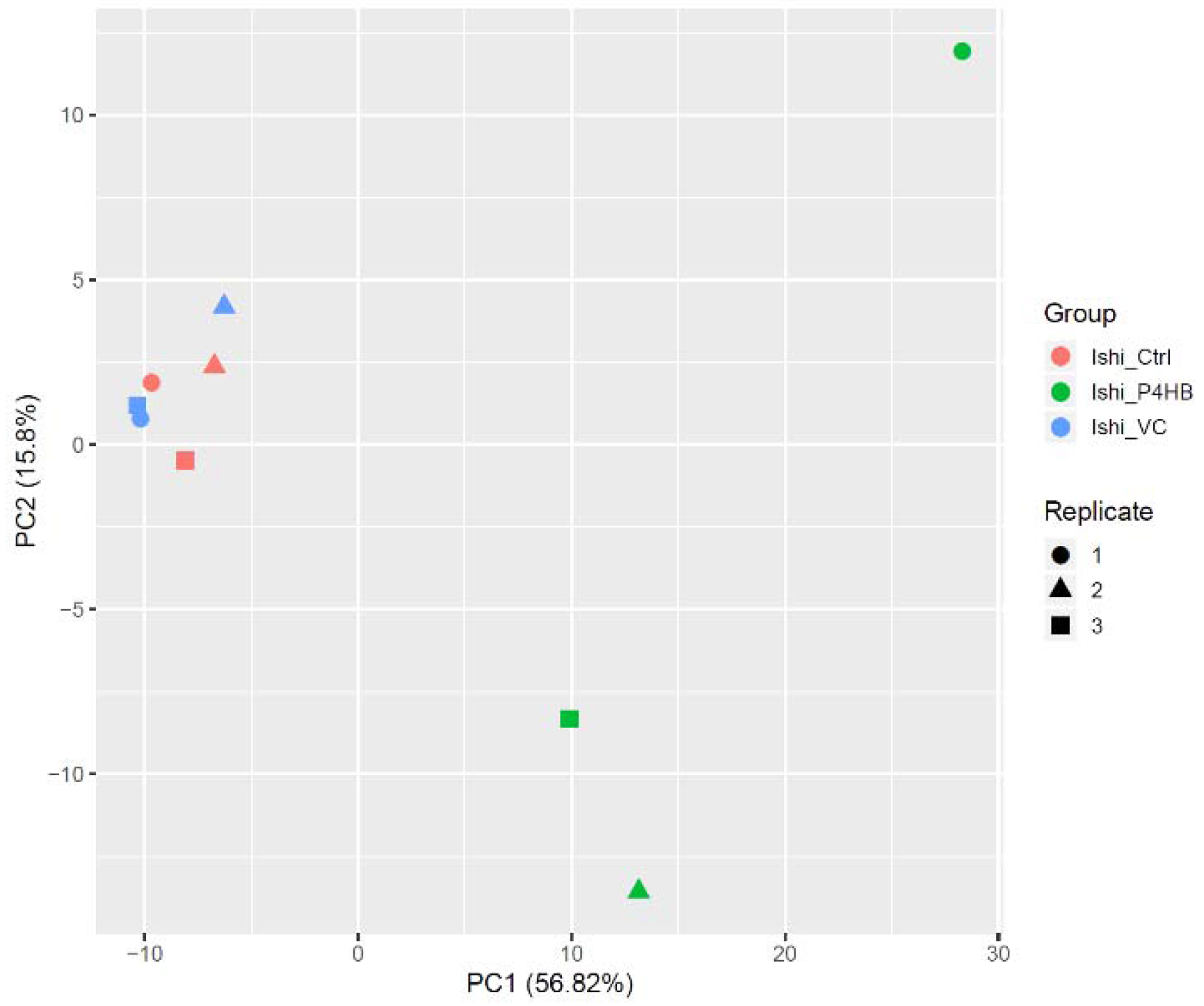
Principal component analysis describing the transcriptional response to rbPDI treatment in human endometrial Ishikawa cells. Control samples (_Ctrl), vehicle control samples (_VC), and rbPDI (_PDI).

**Figure 19.**
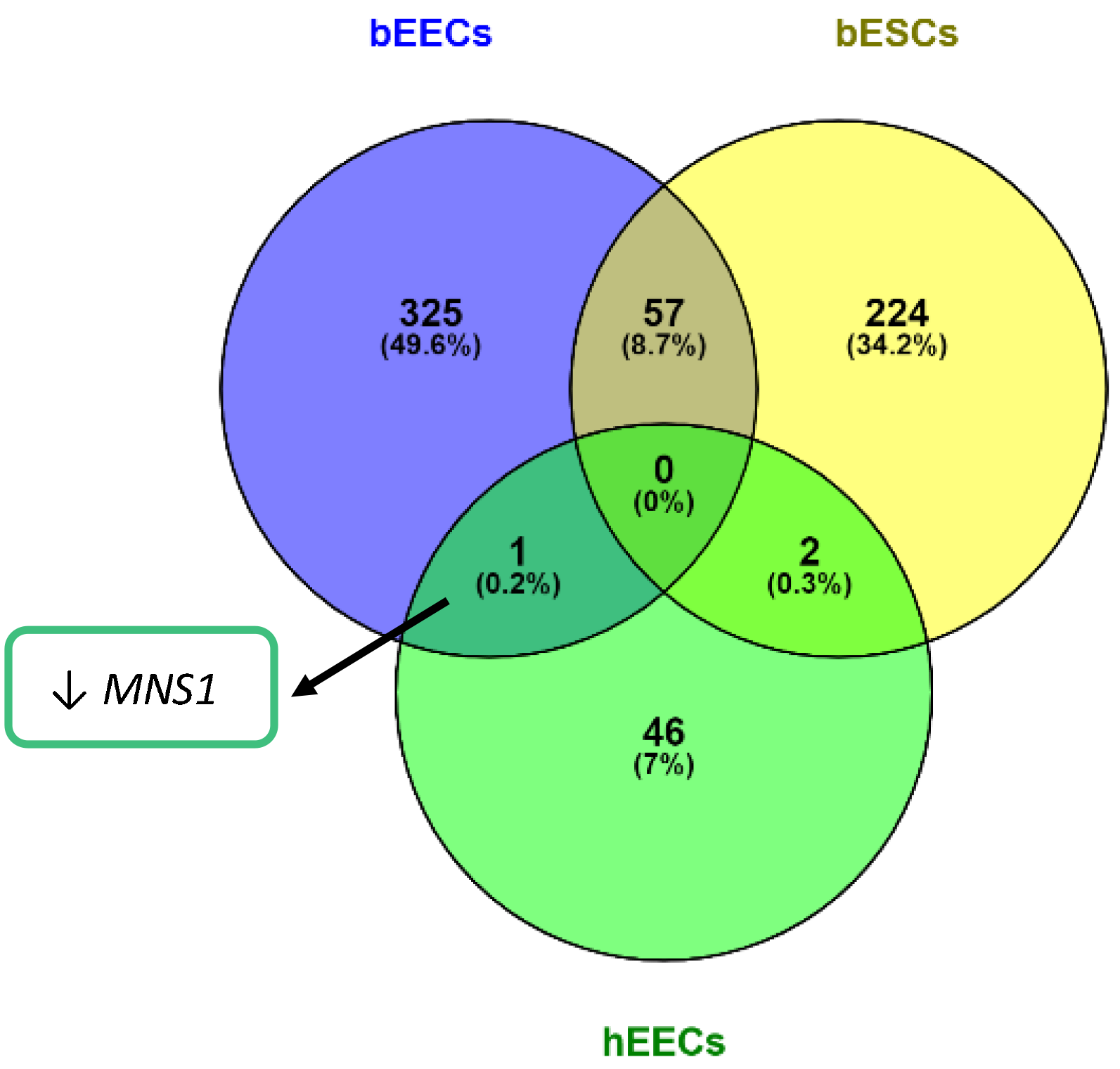
Venn diagram analysis of rbPDI induced DEGs. rbPDI compared to vehicle controls samples (n=3) in human endometrial epithelial cells (hEECs), bovine endometrial epithelial cells (bEECs), and bovine endometrial stromal cells (bESCs). Includes characterised transcripts only. Full data in Supplementary table 21

### *MNS1* knockdown alters *in vitro* trophoblast spheroid attachment to hEECs

Human endometrial cells pre-treated with rbPDI for 48 hours had no significant difference in BeWo spheroid attachment to hEECs compared to those pre-treated with vehicle control at the concentration tested (1000ng/mL) (Figure 20). *MNS1* was commonly downregulated in both hEECs and bEECs in response to rbPDI treatment (Supplementary tables S1 and S18). *MNS1* knockdown was quantified to ensure knockdown of expression to >80%. qRT-PCR demonstrated that *MNS1* was knocked down by 86.1% compared to non-targeting siRNA (Supplementary figure S1). Human endometrial cells pre-treated with siRNA targeting *MNS1* under optimised conditions for 48 hours significantly reduced BeWo spheroid attachment to hEECs when compared to those pre-treated with a non-targeting siRNA (Figure 20).

**Figure 20.**
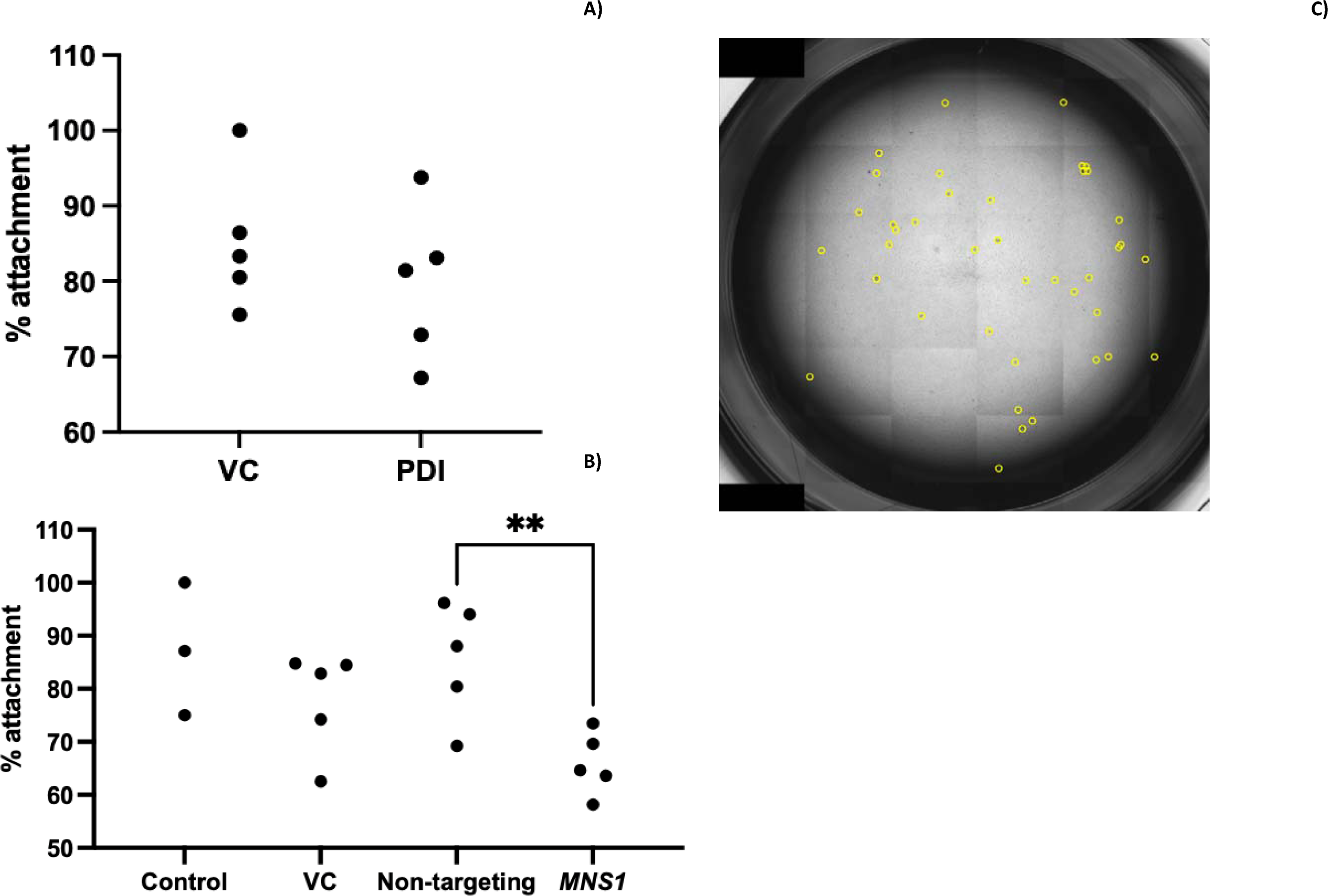
**rbPDI and *MNS1* knockdown effect upon trophobl**ast spheroid **attachment to endometrial cells.** Endometrial epithelial human Ishikawa cells treated with **A)** rbPDI (PDI) or vehicle control (VC), or **B)** siRNA targeting MNS1 or a non-targeting siRNA, were co-cultured with BeWo human trophoblast spheroids to investigate how treatment alters the percentage of spheroids adhering to the endometrial cells (n=5). A) Students t-test or B) ANOVA analysis was used in Graphpad Pr sm to determine significance. Graphs created in Graphpad Prism. OptiMEM only control, lipofectamine only vehicle control. **C)** Representative image of the spheroid attachment assay. Image taken on an EVOS microscope and spheroids counted in QuPath.

### Using a microfluidic device, exposure of endometrial epithelia cells to PDI alters the endometrial secretome in a species-specific manner

Seventeen proteins were identified as differentially abundant within conditioned medium between vehicle control and rbPDI treated hEECs in microfluidic devices (p<0.05) (Table 4). As expected, the protein with the highest fold change is PDI, which was added to the medium. Only 2 of the differentially expressed proteins were identified as human, mitochondrial import inner membrane translocase subunit TIM16 and limbic system-associated membrane protein, the remainder all being identified as bovine proteins.

**Table 4.**
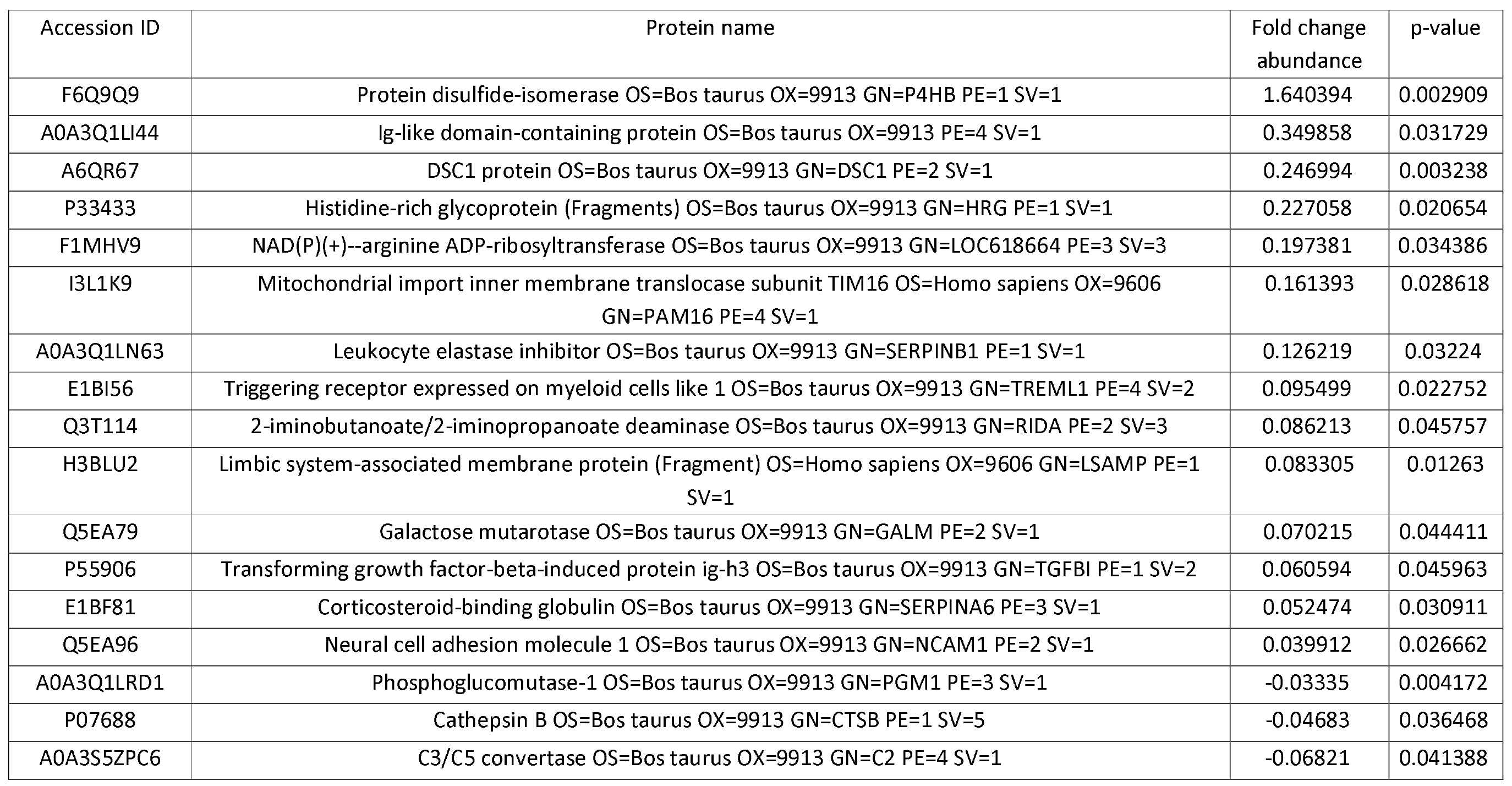
Differentially abundant proteins present in conditioned medium following exposure of human endometrial epithelial cel ls to rbPDI. Proteins identified by tandem mass spectrometry (FDR <0.05), fold change abundance and p-value calculated in excel by students t-test compared to vehicle control samples.

Fifteen proteins were identified as differentially abundant within conditioned medium between vehicle control and rbPDI treated bEECs in microfluidic devices (p<0.05) (Table 5). As expected, bovine PDI was present in the list of differentially expressed genes as it was added to the culture medium. Aside from PDI, 8 proteins: serum amyloid A, fibronectin, tubulin alpha 1D chain, ezrin, alpha-actinin-4, metalloendopeptidase, similar to peptidoglycan recognition protein, and calpain inhibitor were increased in abundance compared to VC, and 6 proteins: collagen type V alpha 1 chain, fibrinogen alpha chain, inter-alpha-trypsin inhibitor heavy chain H1, inter-alpha-trypsin inhibitor heavy chain H4, alpha-2-antiplasmin, and glutathione-independent PGD synthase decreased in abundance compared to VC. Interestingly, serum amyloid A protein was even more highly differentially abundant compared to VC than rbPDI (which was added artificially at a relatively high concentration).

**Table 5.**
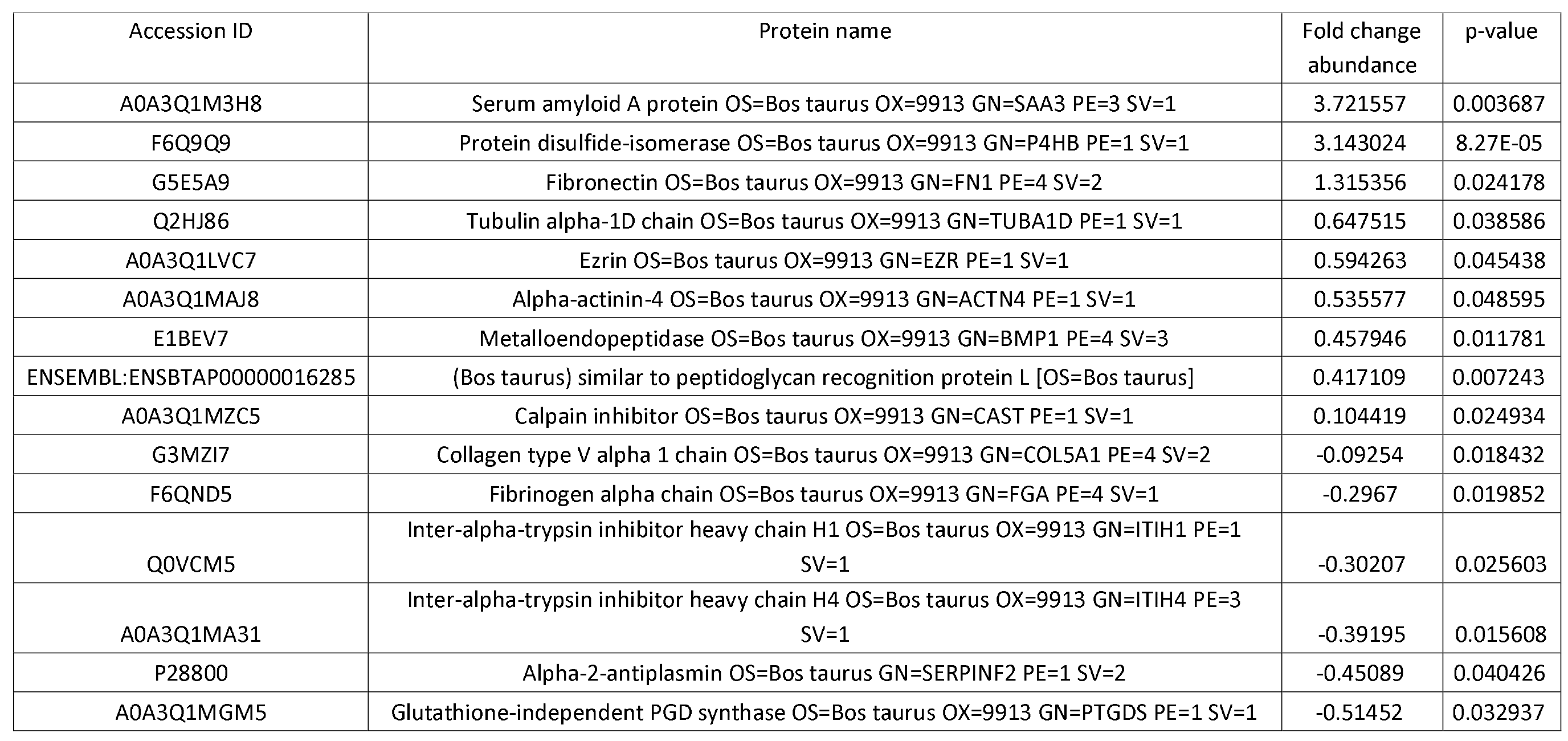
Differentially abundant proteins present in conditioned medium following exposure of bovine endometrial epithelial cel ls to rbPDI. Proteins identified by tandem mass spectrometry (FDR <0.05), fold change abundance and p-value calculated in excel by students t-test compared to vehicle control samples.

## DISCUSSION

We have shown that the PDI protein sequence and function are highly conserved across placental mammals, with rbPDI inducing a significant transcriptional response in bovine and human endometrial cells *in vitro*. Many of the transcripts induced by rbPDI are also induced by roIFNT *in vitro.* The response is cell specific in the bovine, while in human epithelial cells rbPDI induces a limited transcriptional response. One transcript (*MNS1*) was commonly downregulated in both bovine and human endometrial epithelial cells and knockdown of this transcript altered the ability of trophoblast spheroids to attach. rbPDI treatment of bEECs and hEECs under flow in a microfluidic device revealed that rbPDI influences the secretome of endometrial epithelial cells in a species-specific manner.

### PDI secretion supports the transcriptional response to IFNT in the bovine endometrium

The transcriptional response to PDI mostly overlaps with the transcriptional response *in vitro* to roIFNT. Of those DEGs commonly altered by rbPDI and roIFNT when treated separately and in combination when compared to VC, the GO terms antigen processing and presentation, innate immune response and immune effector processes are amongst the highest enriched. These data support the hypothesis that PDI may be involved in the interferon response and/or immune response induced by IFNT. Therefore, this previously uncharacterized embryo-derived protein, PDI, may be secreted by the bovine conceptus to support the actions of IFNT to modulate the endometrial transcriptome or mediate the immune response to IFNT. PDI has previously been linked to immune regulation and could induce tolerogenic dendritic immune cells (35), although the mechanism was not elucidated. We hypothesise that PDI may support the immune modulation effects of IFNT in the bovine endometrium, and may act to tolerise the maternal immune response to the semi-allographic conceptus (36). The KEGG pathways ‘allograft rejection’ and ‘antigen processing and presentation’ were enriched when treated with rbPDI compared to VC, further supporting this hypothesis.

### PDI elicits a unique transcriptional response to IFNT in the bovine endometrium

These data provide further evidence that not all of the transcriptional effects seen within the bovine endometrium *in vivo* in response to the day 15/18 conceptus are due to interferon stimulation (7, 37). Our data supports the hypothesis that the conceptus-derived protein PDI, may be responsible for some of these observed differences (8). As described, there were several DEGs elicited in response to rbPDI alone or in combination with roIFNT, that were not altered by roIFNT alone. This indicates that PDI elicits a unique transcriptional response in the endometrium which may support early pregnancy processes. The transcriptional response to rbPDI in bEECs included multiple enriched GO terms involved in metabolic processes, peptide secretion, and import into cell, which are not enriched in response to roIFNT treatment. Prior to placenta formation, the conceptus relies on the histotrophic secretion from the endometrium into the ULF for nutrients for growth (6). rbPDI may alter metabolic processes, secretion/import in bEECs to optimise production of metabolites for secretion into the uterine luminal fluid (ULF) for conceptus nutrition (38). rbPDI treatment of bEECs also led to the enriched KEGG pathway of ‘cell-adhesion molecules’, which indicates that PDI may be involved in the process of attachment during the initiation of implantation. All the DEGs in response to rbPDI in bEECs mapped to the KEGG pathway ‘cell adhesion molecules’ were upregulated compared to VC, indicating that PDI secretion by the conceptus would increase adhesion to endometrium.

When looking in detail at the DEGs specific to rbPDI or rbPDI+roIFNT treatment compared to VC, but not those induced by roIFNT (22), the GO terms enriched include cell adhesion molecules, metabolic processes, and other immune-related gene sets. Enriched KEGG pathways from this set of DEGS are all related to the immune response, including: NFKB, TNF, IL-17, and toll-like receptor signalling pathways- which are all pro-inflammatory. Although it seems counterintuitive for the conceptus to secrete PDI, which induces a proinflammatory response as seen in this project, studies have shown increases in natural killer cells (39), macrophages (40), or dendritic cells (41) in the bovine endometrium during pregnancy, which are all immune cells recruited by pro-inflammatory signals. PDI may therefore have a role in recruiting these cells to the endometrium, although more evidence is required to confirm this. Overall, these data further support our hypothesis that PDI may support the maternal immune system modulation elicited by IFNT, and potentially facilitate implantation and/or the histotroph secreted by the endometrium.

In bovine endometrial cells *in vitro,* the number of DEGs induced by rbPDI are fewer in stromal than epithelial cells. This may be due to the *in vivo* endometrial architecture, where stromal cells reside below the epithelial layer which lines the uterine cavity. The epithelial cells are therefore the first contact for embryo-derived factors, and these data indicate that the two cell types may respond differently to embryo-derived proteins *in vivo.* Like bEECs, the response to rbPDI in bESCs was associated with GO terms related to immune function and cell-cell adhesion.

### Human endometrial cells have a limited response to PDI *in vitro*

As described, rbPDI altered the expression of fewer transcripts in human endometrial epithelial cells than in bovine endometrial cells. This could indicate a species-specific response to PDI protein, or that the rbPDI transcript interacts differently with human cells than human PDI. Recently published work demonstrated that human endometrial cells responded to recombinant human PDI (with increased expression of microRNA-324-5p), whereas rbPDI induced a different response in bovine endometrial cells (decreased expression of microRNAs -185-5p, -542-3p, and -151a-3p), supporting a model of species-specific response (21). The endometrial transcriptional response in hEECs to rbPDI mostly differed when compared to bEECs, however, the transcript *MNS1* was commonly downregulated in both species in response to rbPDI compared to VC.

*MNS1* encodes the meiosis specific nuclear structural protein (MNS1) which is involved in both cilia and flagella assembly (42). MNS1 has been implicated to be critical in the process of spermatogenesis (43), and extensively researched in the context of male fertility. In endometrium, a recent study described *MNS1* expression being altered in human endometrial organoids in response to oestrogen treatment (44), and in cattle that *MNS1* expression in peripheral white blood cells was significantly different between artificially inseminated compared to natural breeding pregnant heifers (45). The role and mechanism of action of *MNS1* in the endometrium are not currently known and warrant further investigation.

### PDI may mediate implantation at the attachment stage via *MNS1*

Although rbPDI treatment of human endometrial Ishikawa cells had no significant effect on BeWo trophoblast cell spheroid attachment in this project, recent work demonstrated that inhibiting PDI significantly increases attachment of Jeg-3 spheroids onto AN3CA human endometrial cells (18). Interestingly, PDI expression was found to be significantly higher in AN3CA cells compared to Ishikawa cells, and Ishikawa cells are considered to be in a ‘receptive’ state (46) whereas AN3CA cells are not (47). One possible explanation is that PDI acts to decrease attachment to non-receptive endometrium, acting as a timing mechanism to ensure attachment only to a receptive endometrium or even acting as a mechanism to prevent premature implantation, such as in the fallopian tube. One study supporting this hypothesis demonstrated that PDI expression is significantly higher in the stromal cells of non-receptive endometrium of bonnet monkeys compared to receptive endometrium (48). Although the study also found no difference in PDI expression in the epithelial glands, the authors do not appear to have assessed the luminal epithelium.

Although implantation attachment wasn’t shown to be altered by rbPDI in this investigation, there are limitations to the work done here. rbPDI may not function in exactly the same way as the human ortholog, and therefore may not act upon the human endometrial Ishikawa cells to produce an effect on attachment. Also, only one concentration of rbPDI was tested, whereas influencing trophoblast spheroid attachment may occur at a specific range of concentration. When *MNS1* expression was decreased by >86% in Ishikawa cells, the rate of BeWo spheroid attachment significantly decreased compared to controls. *MNS1* expression was decreased in response to rbPDI treatment in both hEECs and bEECs. Therefore, exposure to PDI may decrease trophoblast attachment to the endometrium, supported by work demonstrating that PDI inhibition increases spheroid attachment (18), and that when *MNS1* expression is reduced (as occurs in PDI exposure) attachment reduces. This work shows that *MNS1* may be a mechanism by which PDI mediates implantation, perhaps in a reproductive cycle timing specific manner.

### rbPDI alters the endometrial secretome in a species-specific manner

Collection of the conditioned or ‘spent’ medium from the microfluidics devices seeded by hEECs or bEECs allowed the secretome of the cells to be analysed by mass spectrometry. Assigning protein IDs to the proteins secreted by hEECs showed that only 2/17 proteins which were differentially abundant were assigned homo sapiens species identifiers. This discrepancy is speculated to be due to the bovine FBS used in the culture medium and/or false positives. The 2 human differentially abundant proteins, mitochondrial import inner membrane translocase subunit TIM16 and limbic system-associated membrane protein (fragment), were only slightly higher in abundance compared to VC (0.65- and 0.08-fold change respectively) in the conditioned medium. TIM16 was in 65% greater abundance in the conditioned medium after rbPDI treatment than VC, and is involved in ATP-dependant protein translocation into the mitochondrial matrix as a component of the TIM23 complex (49). TIM23 is involved in the import of approximately 60% of mitochondrial proteins, and alterations in the mitochondrial proteome is associated with changes in growth conditions and are mostly involved in energy metabolism (50). Therefore, the secretion of TIM16 by endometrial cells in response to PDI could aid the function of the conceptus mitochondria to increase energy availability and growth, and as a result increased viability of the conceptus.

There were 0 commonly differentially abundant proteins present in the conditioned medium between hEECs and bEECs when compared to vehicle control samples. Therefore, the response to rbPDI altering the secretome of these cells may be species-specific.

In bEEC conditioned medium serum amyloid A protein was the most highly differentially abundant protein, at 3.72-fold the amount of serum amyloid A present following treatment with rbPDI compared to control. Serum amyloid A is involved in cell-cell communication (51), but was not found in the ULF from pregnant cattle on day 15, 16 or 18 (8, 52), indicating that it may be an artifact of *in vitro* cell culture treatment with rbPDI, or that it is only produced very locally and/or immediately utilised *in vivo*. Tubulin alpha 1D chain has however been identified in pregnant ULF on day 16 from cattle (8, 52, 53) and was increased 0.64-fold during rbPDI treatment. Interestingly, tubulin alpha 1D was also secreted by bovine day 16 conceptuses *in vitro* (53).

In conclusion we have demonstrated that PDI, a conceptus-derived protein, with highly conserved amino acid sequence across placental mammals alters the transcriptome of the endometrial epithelium in a species-specific manner. In bovine PDI enhances the IFNT simulated response during the peri-implantation period of pregnancy, as well as modifying PDI-specific transcripts in the endometrial epithelial and stromal cells. One transcript, *MNS1* is modified by PDI in two species with divergent implantation strategies i.e., human and bovine, and modifies the ability of BeWoW spheroids to attach. Exposure of epithelial cells to PDI also modified the secretome in a species-specific manner. Collectively these data demonstrate that PDI plays a species-specific role in endometrial function in mammals with different implantation strategies.

## ACKNOWLEDGEMENTS

Research in NF’s lab is supported by N8 agri-food pump priming, QR GCRF, as well as BBSRC grant numbers BB/R017522/1, as well as support from LTHT. NF and MO’C are funded jointly by BBSRC grant BB/X007332/1 and Wellcome Trust grant 227178/Z/23/Z.

---

**Supplementary Figure 1.**
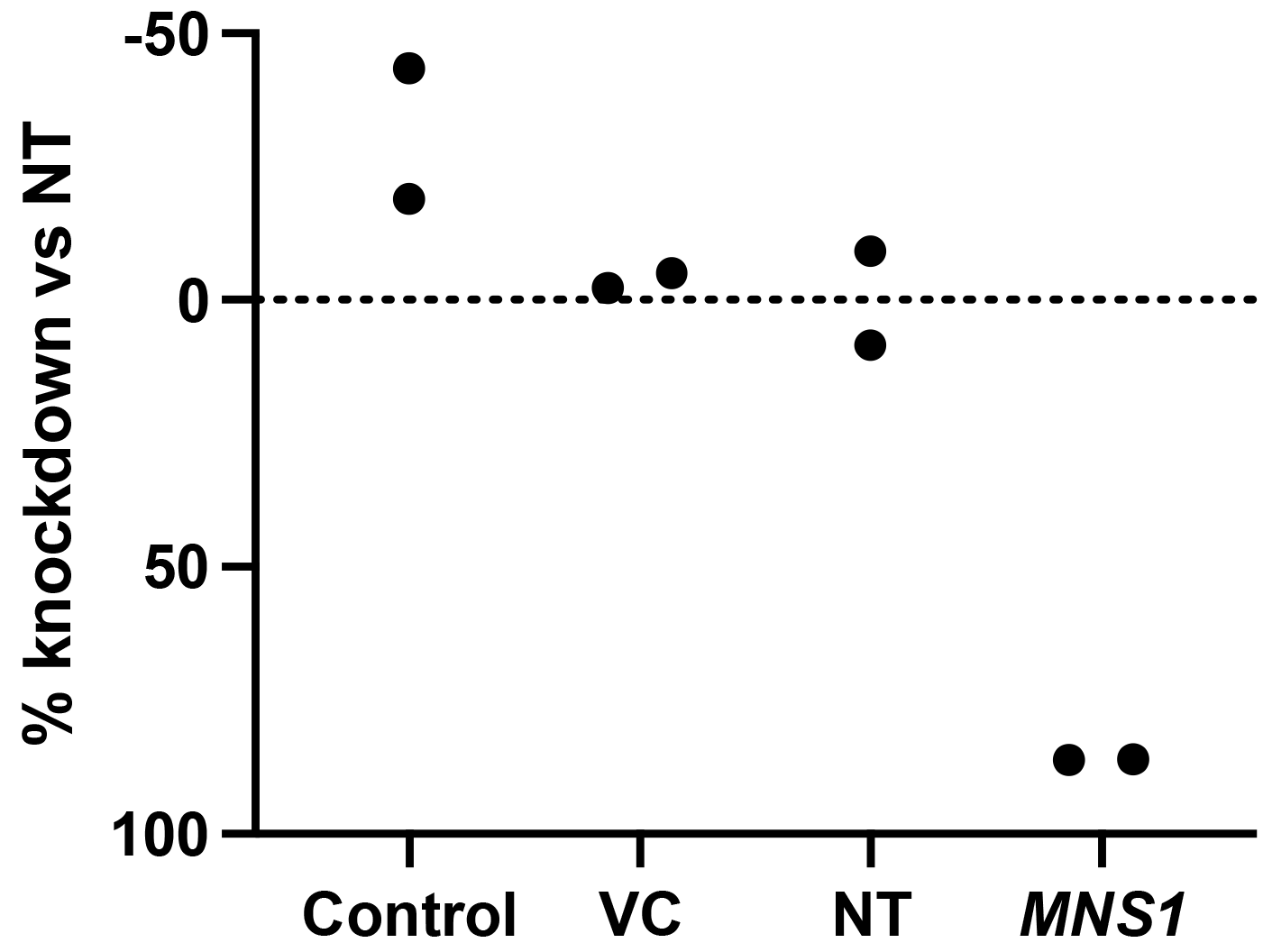
siRNA knockdown efficiency *MNS1* compared to non-targeting siRNA control (NT). OptiMEM only media control, lipofectamine vehicle control (VC), non-targeting siRNA (NT), and siRNA targeting *MNS1* treated Ishikawa cells (n=2) for 48 hours. *MNS1* expression determined by qRT-PCR and % knockdown calculated from the 2^-ΔΔCt^ values using *ACTB* as a normaliser gene relative to the non-targeting siRNA treated samples. Figure created in Graphpad Prism.

